# ERV-derived dsRNAs induce stress granules that drive neural crest fate to initiate melanoma

**DOI:** 10.64898/2026.11.06.764901

**Authors:** Kyle D. Drake, Emily Formato, Siras Hakobyan, Rachel A. Hongo, Song Yang, Jennifer Porat, Owen Sanborn, Anne L. Robertson, Dmitry A. Kretov, Yi Zhou, Daniel Cifuentes, Manfred Kunz, Hans Binder, Ryan A. Flynn, Leonard I. Zon

## Abstract

Melanoma arises when melanocytes are oncogenically reprogrammed into neural crest-like cells, but what initiates this switch is unknown. Here we show that endogenous retrovirus (ERV)-derived double stranded RNAs (dsRNAs) induce stress granules (SGs) that drive this fate change. In zebrafish, melanoma progresses through a stage-specific ERV switch: *zferv* marks precursor cells before *crestin* activates in tumor cells. Both elements generate dsRNA that triggers *eif2ak2*-dependent SGs within precursor lesions and through transformation. SGs are tumorigenic as *G3BP1* overexpression accelerates melanoma onset, whereas mosaic, melanocyte-specific *g3bp1/2* knockout delays it. Small molecule-mediated inhibition of G3BP1/2 eliminates precursor lesions and reduces tumor burden. Similarly, blocking dsRNA sensing prevents SG formation and lowers melanoma incidence. Human nevi and melanomas show analogous, stage-specific ERV programs and SG assembly. In human melanoma cells, SGs sequester and translationally repress SOX10-regulating mRNAs, including the tumor suppressor *KMT2D*. Mosaic knockout of *kmt2d in vivo* enhances SOX10-driven neural crest reactivation. Thus, an ERV-SG axis establishes an oncogenic translatome that enables melanoma fate, with diagnostic and therapeutic potential.

## Introduction

In many cancers, the initiating cell of a tumor reprograms into its embryonic, progenitor-like state prior to overt tumor formation^1^. In melanoma, the melanocyte de-differentiates into a neural crest-like state during initiation^2^. This phenomenon can be observed in zebrafish as reactivated expression of a retroelement *Gag* gene, *crestin*, which is normally expressed only in neural crest cells during embryogenesis^3–5^. We recently identified a melanoma precursor lesion called the cancer precursor zone (CPZ), in which oncogenically transformed, *mitfa:hBRAF^V600E^;p53^-/-^*melanocytes display *mitfa* upregulation prior to transformation into the neural crest-like state^6^. Whereas not all CPZs form tumors, virtually all *crestin+* lesions give rise to melanoma^3,6^. This observation demonstrates that molecular events beyond primary oncogenic insults are required to induce a full, cancer - competent fate shift.

*Crestin* reactivation in early melanomas suggests that dysregulation of *bhikhari-2 (bhik2)* endogenous retroviruses (ERVs) occurs *in vivo* during melanoma initiation. ERV reactivation is an emerging hallmark of cancer biology and may be tumorigenic^7^. Upregulation of ERVs in human melanoma, such as *HERV-K*, is associated with malignant transformation and corresponds with cancer stemness^8–10^. Analogously, expression of the *crestin bhik2-LTR*, which encodes a double stranded RNA (dsRNA), accelerates melanoma onset in zebrafish^11^. The same phenotype is observed with overexpression of human ERVs^11^. This indicates a shared oncogenic role for viral-like dsRNA substrates in tumor initiation, although the mechanism by which this occurs remains unclear.

One molecular outcome of intracellular dsRNA detection is stress granule (SG) formation^12^. SGs are cytoplasmic, ribonucleoprotein condensates formed upon stalled translation initiation^13^. Various stressors, including temperature stress, oxidative stress, and viral stress can induce SG formation^14,15^. As SGs are cytoprotective, they are leveraged by cancer cells to withstand the harsh biological environments associated with tumorigenesis^16–18^. SGs have also been shown to enhance translation of pluripotency factors governing tumor-initiating cell identity^19^. To date, a tumorigenic role for SGs has been implicated in acute myeloid leukemia, breast cancer, cervical cancer, colorectal cancer, gastric cancer, glioblastoma, medulloblastoma, melanoma, non-small cell lung cancer, oral cancer, osteosarcoma, ovarian cancer, pancreatic ductal adenocarcinoma, prostate cancer, and renal cell carcinoma^17^. Despite these findings, the role of SGs in an endogenous, *in vivo* tumor model remains unexplored. We hypothesized that ERV expression during melanoma initiation induces SGs that subsequently impart an oncogenic translatome to facilitate acquisition of neural crest fate.

Here we use a primary zebrafish melanoma model where melanocytes express human *BRAF^V600E^* under control of the *mitfa* promoter and lack the tumor suppressor *p53*^20^. These melanocytes develop a cancerized field (CF) of oncogenically competent melanocytes^6^. Some CF melanocytes form a CPZ, and a subset of CPZs transform into *crestin*+ melanomas^3,6^. Using this model, we show that stage-specific ERV-derived dsRNAs trigger SG formation beginning in CPZs and continuing in melanomas. We identify a similar, stage-specific ERV program defining human nevi from human melanoma and uncover SG formation and persistence in human melanocytic neoplasms. Mechanistically, we show that SGs impart an oncogenic, neural-crest priming translatome upon sequestration of mRNAs encoding tumor suppressive epigenetic factors centered around SOX10 regulation. We use *kmt2d* as an exemplar to demonstrate that downregulation of SG-controlled mRNAs can enhance the ability of SOX10 to induce the neural crest program as read out through chromatin profiling and RNAseq. Disrupting the core SG regulators *g3bp1/2* through genetic and chemical approaches, or abrogating SG formation by limiting eif2ak2-mediated dsRNA sensing, reduces melanoma incidence and overall tumor burden. This positions ERV-dysregulation and SG biology as diagnostic tools and therapeutic vulnerabilities that can be readily exploited to halt cancer initiation and progression.

## Results

### Stage-specific expression of stress granule-inducing ERVs occurs during zebrafish melanomagenesis

We employed our model of zebrafish melanoma to understand ERV expression dynamics across cancer initiation. In our system, *mitfa*:*BRAF^V600E^;p53^-/-^* melanocytes comprise a cancerized field (CF), of which a subset upregulate *mitfa* and develop a cancer precursor zone (CPZ)^6^ (Fig. 1A). The CPZ is a precancerous lesion in which a subset of cells reprogram into a neural crest-like state, as defined by their reactivated expression of the embryonic, *bhik2* ERV *crestin*^6^ (Fig. 1A). Each stage of melanoma initiation has increased malignant potential, with *crestin*+ patches representing the earliest *bona fide* tumor cells^6^. We first sought to explore whether ERV expression in the CPZ stage was distinct from early melanoma cells. TEtranscripts-mediated analysis of transposable element (TE) expression using bulk RNAseq data from CPZs and early tumors revealed a substantial number of retroelements, including ERVs/LTRs, overrepresented in the CPZ stage (Fig. 1B; Supp. Fig. 1A). While total ERV/LTR counts do not significantly change during the CPZ to melanoma transition, we noticed the major ERV/LTR expressed in CPZs or melanomas was distinct (Fig. 1B-C, Supp. Fig. 1B-C). The internal region of *zferv* was found to spike in expression during the CF to CPZ transition then undergo significantly decreased expression in the tumor stage as *bhik2* emerges (Fig. 1B-C; Supp. Fig. 1D). The *zferv-LTR* was mildly elevated from CF to CPZ and similarly downregulated in tumors (Fig. 1B-C; Supp. Fig. 1E). This stage-specific ERV expression switch was validated using a second TE-quantification algorithm, SQuIRE (Supp. Fig. 1D-F). *zferv* is a zebrafish-specific ERV distinct from *bhik2* elements. There are at least eight *zferv* pseudogenes, with two encoding full proviruses and six that are degenerate^21^. Stage-specific ERV expression dynamics were corroborated using CUT&RUN for H3K27ac, which showed a doubling in H3K27ac abundance exclusively at the CPZ stage for both the internal region and LTR elements of *zferv*, whereas *bhik2* elements gained H3K27ac only in tumor cells (Fig. 1D-E). As upregulation of *mitfa* is the marker for CPZ onset, we explored whether *zferv* loci contain MITF binding sites^6^. Nearly all *zferv* loci were found to contain significant MITF motifs, with a degenerate *zferv* at chr19 having the most mRNA expression by RNAseq, largest H3K27ac peak, and strongest predicted MITF binding site motif (Fig. 1E; Supp. Fig. 1G-H). This data demonstrates that stage-specific ERV programs accompany zebrafish melanomagenesis.

**Figure 1.**
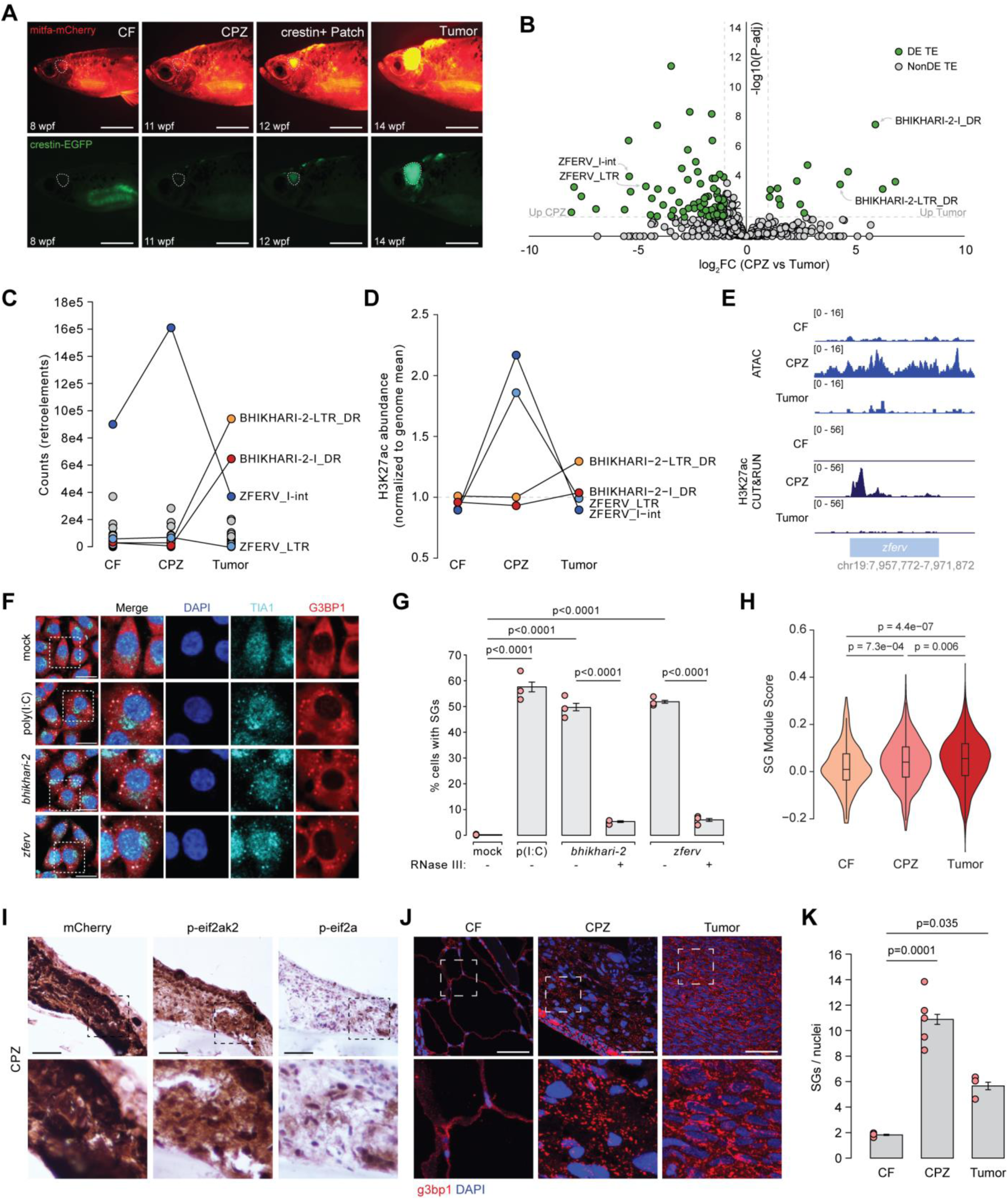
Stress granules form during zebrafish melanomagenesis upon ERV stress. (**A**) Fluorescent live imaging of zebrafish melanoma development. White outline region shows cancerized field (CF) that develops into a cancer precursor zone (CPZ; mitfa-mCherry^high^) prior to crestin-eGFP activation (early tumor patch that sprouts overt tumor). wpf=weeks post fertilization. (**B**) Volcano plot of transposable element (TE) expression in CPZ (n=4) vs tumor (n=3). Green dots are significantly differentially expressed (DE). (**C**) Retroelement expression across melanoma stages showing predominant expression of *zferv* in CPZs and *bhikhari-2* in tumors. (**D**) Line plot showing abundance of H3K27ac at *zferv* and *bhikhari-2* elements across melanoma initiation upon CUT&RUN for n=6 CF (pooled n=1), n=6 CPZ (pooled n=1), and n=2 tumors (average value shown). (**E**) IGV tracks showing ATAC (top) and H3K27ac CUT&RUN (bottom) at *zferv* chr19 locus in CF, CPZ, and tumor samples. (**F**) IF for G3BP1 (red) and TIA1 (cyan) using A375s lipofected with dsRNA mimic poly(I:C), *bhikhari-2* dsRNA, or *zferv* dsRNA for 2 hours. (**G**) Bar chart showing quantification of percent of cells with SGs when lipofected with dsRNAs (n=3). Significance determined via ANOVA with post-hoc Tukey’s test. (**H**) Violin plot showing Seurat AddModuleScore feature expression of core SG genes across melanoma stages. Significance determined by Wilcoxon rank-sum test. (**I**) Immunohistochemistry for mCherry, p-eif2ak2, and p-eif2a in representative CPZ. (**J**) IF for g3bp1 (red) with DAPI nuclear counterstain (blue) in representative CF, CPZ, and tumor sections. (**K**) Quantification of stress granule (SG) abundance in CF (n=3), CPZ (n=5), and tumor (n=3). Error bars represent SEM. Significance determined by ANOVA with post-hoc Tukey’s test. Scale bars, 0.5 cm (**A**), 25 µm (**F**), 500 µm (**I**), 50 µm (**J**).

Overexpression of both human and zebrafish ERVs are tumorigenic in melanoma^11^. Since these elements are not orthologous, their shared tumorigenicity implicates a conserved downstream mechanism rather than element-intrinsic functions. ERV expression generates viral-like dsRNAs that activate antiviral pathways including SG formation, which is broadly tumorigenic^4,22–24^. We tested directly whether the stage-defining zebrafish ERVs, *zferv* and *bhik2*, induce SGs when expressed as dsRNAs in cultured human melanoma cells. We found 52% and 50% of cells contained SGs after 2 hours of *zferv* and *bhik2* dsRNA treatment, respectively, which was on par with 58% of cells displaying SGs when treated with the dsRNA mimic poly(I:C) (Fig. 1F-G). SG induction was dsRNA-dependent as this phenotype was abolished with RNase III treatment of the dsRNAs prior to transfection (Fig. 1G; Supp. Fig. 2). Analysis of zebrafish melanoma scRNAseq revealed that core SG genes are markers of oncogenically transformed melanocytes and are progressively upregulated during melanomagenesis (Fig. 1H; Supp. Fig. 3). We therefore predicted that zebrafish melanoma precursor cells and tumor cells would contain SGs *in vivo* upon ERV stress.

For ERV-derived dsRNAs to induce SGs, they first must be sensed by the cytosolic dsRNA-binding protein eif2ak2, which results in its autophosphorylation^25^. We found by western blot that CPZ and tumor cells have significantly increased p-eif2ak2 compared to melanocyte-null and tumor-adjacent skin (Supp. Fig. 4). In turn, autophosphorylation of eif2ak2 can induce phosphorylation of eif2a^26^. Immunostaining CPZs and melanomas indeed revealed strong p-eif2ak2 signal and the induction of p-eif2a, which was unobserved in control melanocytes (Fig. 1I; Supp. Fig. 5A-B). As eif2a is a translation initiation regulator, its phosphorylation results in the accumulation of stalled translation initiation complexes that trigger SG assembly^27^. Immunofluorescence for the central SG nucleator g3bp1 revealed an induction of SGs in CPZs that persist in tumors (Fig. 1J-K; Supp. Fig. 5C). Similar SG condensate staining pattern was observed in CPZs upon immunofluorescence for other core SG markers, including tia1l and the dsRNA-binding protein stau2 (Supp. Fig. 6). This indicates SGs form naturally upon ERV stress during melanomagenesis.

### Human melanoma displays stage-specific ERV programs and tumorigenic stress granule assembly

Predicting which human precursor lesions transform into melanoma remains a clinical challenge^28,29^. We reasoned that a similar *zferv/bhik2* ERV switch might occur in human melanoma, which could ultimately aid diagnostic ability. To test this, we quantified TE expression in 22 nevi and 46 melanomas. These samples have been previously profiled and classified into two major subtypes, differing in genetic background, gene signature, and immune-state, among other parameters^30^. We found that holistic TE expression weakly separated sub-populations of nevi and melanomas along principal component space (Fig. 2A). Of all differentially expressed TEs, most were of the ERV/LTR class (Supp. Fig. 7A). We used the 15 patient-matched nevus-melanoma datasets and found 20 significantly differentially expressed ERVs between nevi and melanoma, with 13 higher in nevi and 7 higher in tumors (Fig. 2B). The expression ratio of these 20 ERVs was upregulated in melanomas vs nevi in all paired samples, independent of subtype (Fig. 2C). We then analyzed this expression ratio in the remaining 38 unpaired samples and found a similar separation of nevi from melanoma with an AUC of 0.99 (Fig. 2D; Supp. Fig. 7B). To test whether this reflected general ERV transcriptional trends, we built randomly drawn ratios of ERV expression matched in number to each side of the signature and found significantly weaker discrimination, with a median AUC of 0.64 (Supp. Fig. 7C). This finding reveals a robust set of ERVs whose expression distinguishes melanoma precursor lesions from tumors.

**Figure 2.**
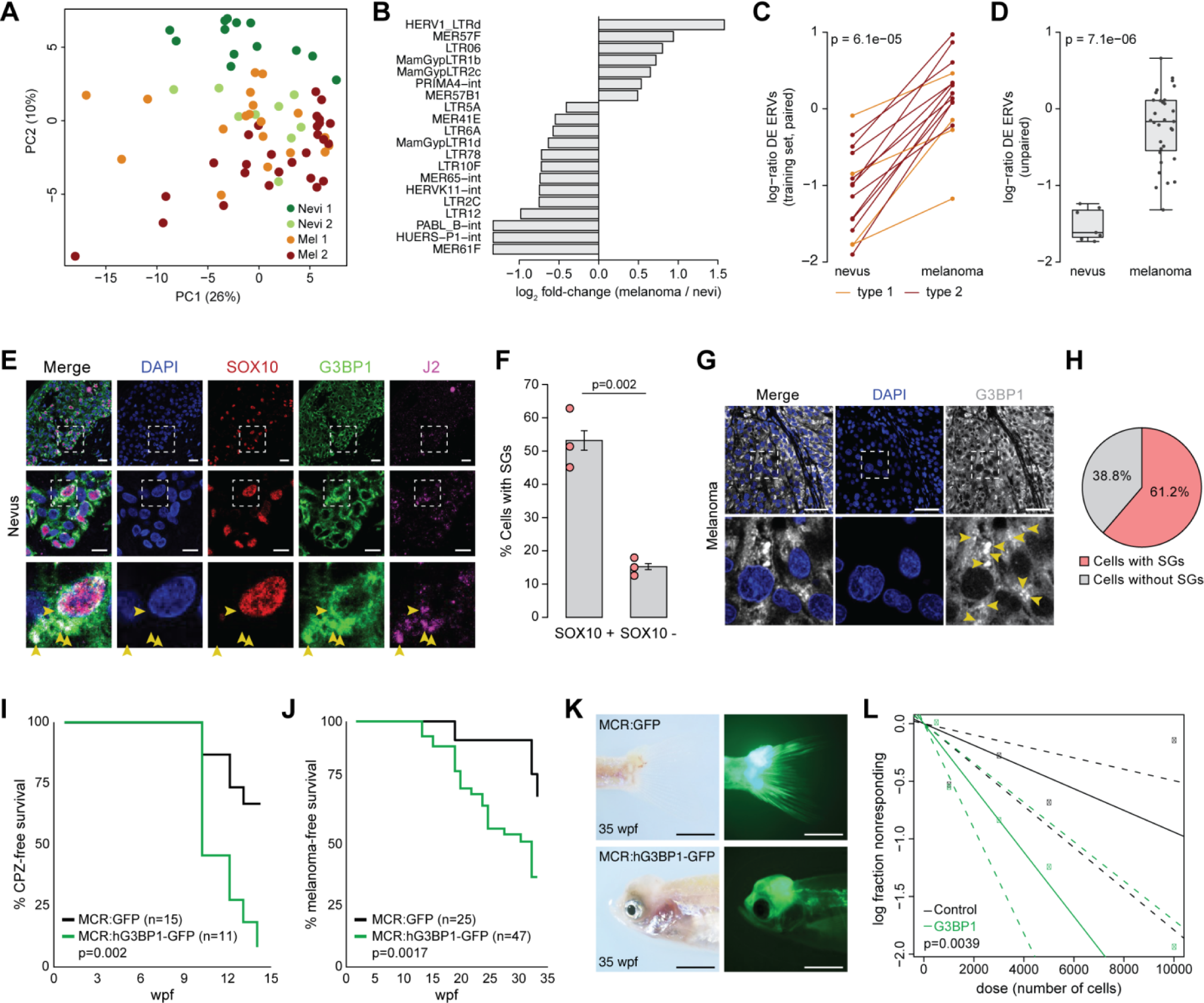
Human stage-specific ERV programs accompany melanomagenic stress granule formation. (**A**) Principal component (PC) analysis of human nevi and melanoma subtypes based on transposable element (TE) expression. (**B**) Bar chart showing log_2_ fold-change for the 20 significantly differentially expressed ERVs/LTRs between human nevi and melanoma. Significance determined by DESeq2 with a paired design (∼ patient + Status); Benjamini-Hochberg-adjusted p < 0.05. (**C**) Line plot showing ratio of the 20 ERVs/LTRs across paired nevi-melanoma datasets. The ratio is higher in the melanoma than the nevus in all 15 pairs. (**D**) Box plot showing the expression ratio of the 20 ERVs/LTRs in unpaired human samples, which were not included in the initial training set. AUC = 0.99; significance determined by label permutation (10,000 iterations). (**E**) IF for SOX10 (red), G3BP1 (green), and dsRNA antibody J2 (magenta) with DAPI nuclear counterstain (blue) in human nevus. Arrowheads point to SGs. (**F**) Quantification of SGs in SOX10+ vs SOX10-cells in human nevi (n=3). Error bars represent SEM. Significance determined by unpaired T-test. (**G**) IF for G3BP1 (white) with DAPI nuclear counterstain (blue) in human melanoma. Arrowheads point to SGs. (**H**) Pie chart showing percent of human melanoma cells that contain SGs (n=2). (**I-J**) Kaplan-Meier curves showing CPZ (**I**) and tumor (**J**) onset rate between *MCR:GFP* and *MCR:hG3BP1-GFP* zebrafish. wpf=weeks post-fertilization. Significance determined by log-rank test. (**K**) Representative image of *MCR:GFP* (top) and *MCR:hG3BP1-GFP* (bottom) zebrafish melanoma validating overexpression of transgene. (**L**) Extreme limiting dilution assay plot for *MCR:GFP* (n=38) and *MCR:hG3BP1-GFP* (n=41) melanoma transplants. Scale bars, 50 µm (**E**, **G**), 0.5 cm (**K**).

In this dataset, *EIF2AK2* was significantly upregulated in melanomas versus nevi (Supp. Fig. 7D). As *EIF2AK2* transcription is induced downstream of dsRNA sensing, the observed upregulation suggested pathway activation, which would predict SG induction as seen in zebrafish^31^. We therefore tested whether human dysplastic lesions and melanoma cells possess SGs by performing IF for G3BP1, SOX10 (to mark melanoma precursors), and J2 (to mark dsRNA) using sections of human nevi. We detected a significant induction of SGs in SOX10+ cells when compared with SOX10-adjacent cells (Fig. 2E-F; Supp. Fig. 8). SGs in SOX10+ cells could be observed to contain dsRNA (Fig. 2E). Immunostaining established tumors showed SG formation in ∼61% of human melanoma cells (Fig. 2G-H). Like zebrafish, core SG genes are upregulated in human melanoma (Supp. Fig. 9A). In particular, *G3BP1* shows a significant, positive transcriptional correlation with *MITF* in melanomas compared to normal skin (Supp. Fig. 9B). Taken together, ERV-induced SG formation is likely to participate in human melanomagenesis.

To test the *in vivo* tumorigenicity of SGs, we cloned human *G3BP1-GFP* into our melanocyte-specific expression vector called MiniCoopR (MCR)^32^. We confirmed in embryos that hG3BP1 does not endogenously induce SGs but readily condenses under stress (Supp. Fig. 10A-C). Longitudinal imaging analysis revealed accelerated CPZ onset and increased CPZ abundance in *MCR:G3BP1-GFP* zebrafish relative to *MCR:GFP* expressing controls (Fig. 2I; Supp. Fig. 10D). In turn, *MCR:G3BP1-GFP* zebrafish had faster melanoma onset and developed more tumors per fish (Fig. 2J-K; Supp. Fig. 10E). *MCR:G3BP1-GFP* tumor cells had significantly greater engraftment potential than *MCR:GFP* controls when transplanted into irradiated casper recipients (Fig. 2L; Supp. Fig. 10F-G). These data demonstrate an evolutionarily conserved, tumorigenic role for SG induction during melanomagenesis.

### KMT2D mRNA sequestration by stress granules facilitates SOX10-driven neural crest reprogramming

We reasoned that the oncogenic impact of persistent SG formation during melanoma would be their ability to sequester mRNA and shift the translational landscape^33^. Since SGs emerge prior to overt tumor formation, we hypothesized that the mRNA regulation exerted by SGs would facilitate neural crest fate acquisition. To identify the mRNAs under SG control in melanoma cells, we performed G3BP1 RIP-seq with concurrent polysome sequencing using human A375 cells stressed with sodium arsenite, a rapid and extremely robust stressor (Fig. 3A; Supp. Fig. 11A-B). Overlap analysis identified 47 mRNAs that were sequestered by G3BP1 and less translated upon SG formation (Fig. 3A-B). SG-regulated mRNAs enriched for epigenetic functions, including H3K4 methylation (Supp. Fig. 11C). This set of epigenetic regulators is likely tumor suppressive as their expression signature is significantly downregulated in human melanoma (Supp. Fig. 11D). In addition, we noticed that some of these genes, such as *FLG* and *KMT2D*, have high mutational burden in human melanomas (Fig. 3C). We assessed the average mutation percentage of genes whose mRNA is regulated by SGs and found ∼2.1x increase in mutational burden compared with the whole genome, indicating SGs sequester mRNAs whose genes are natural targets of genomic alterations (Fig. 3C; Supp. Fig. 11E). These genes are expressed at levels equal to or higher than the general transcriptome, ruling out the possibility that this set reflects lowly expressed genes subject to high mutation burden in melanoma^34^ (Supp. Fig. 11F). This molecular redundancy reveals that SGs are positioned to impose and/or compound loss-of-function events for commonly altered tumor suppressive programs.

**Figure 3.**
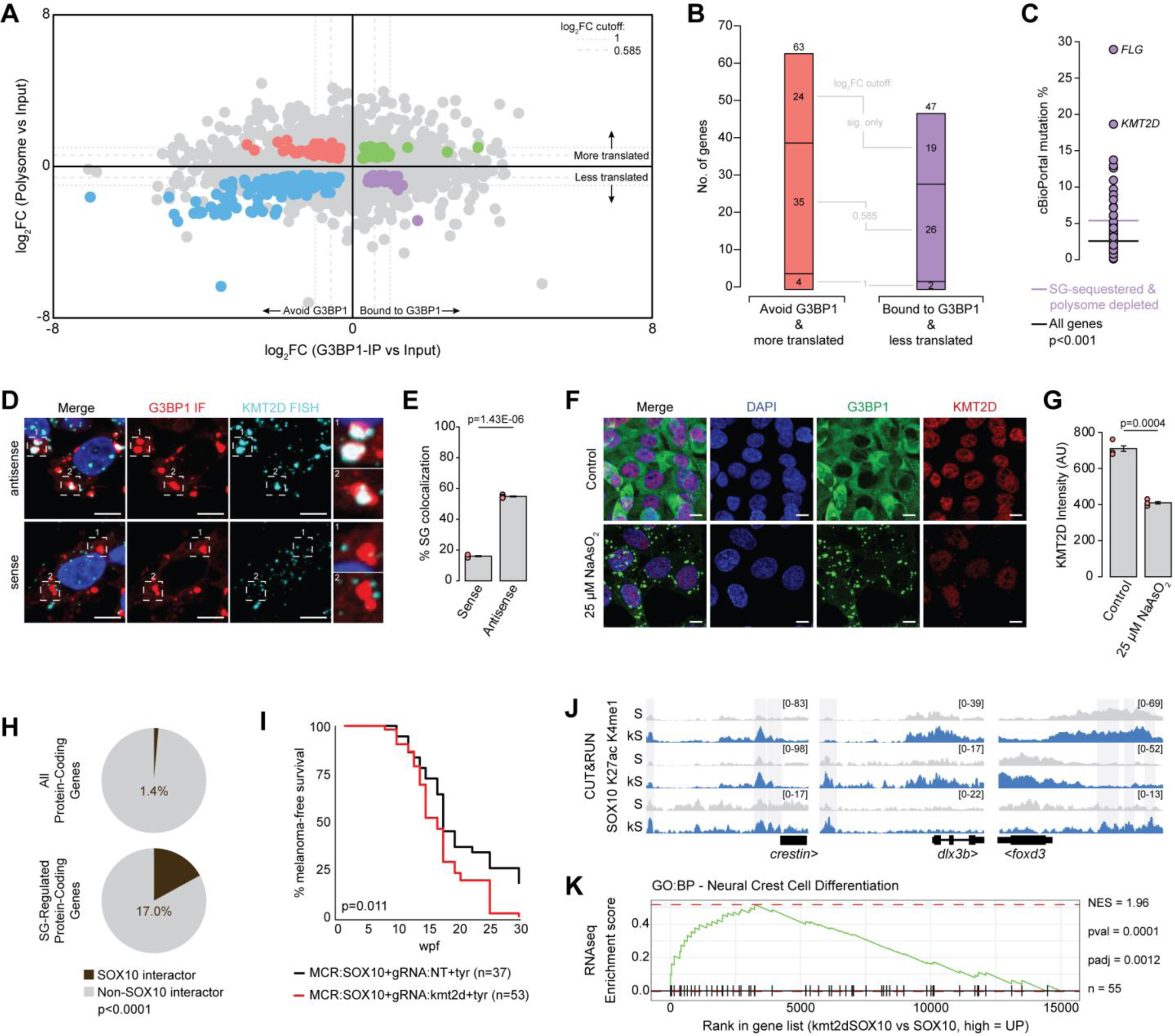
Stress granules capture KMT2D mRNA to facilitate SOX10-driven neural crest fate. (**A**) Cumulative plot showing G3BP1 RIP-seq log_2_FC (X-axis) and polysome-seq log_2_FC (Y-axis) in A375 cells treated with 500 µM sodium arsenite for 1 hour. Colored dots are statistically significant (p-adj<0.05). Different dashed lines show different thresholds of log_2_FC cut-off. (**B**) Bar chart showing number of genes identified to avoid SGs and show preferential translation (red) and captured by SGs with concomitantly reduced translation (purple) per log_2_FC cut-off. (**C**) Mutation percentage for each gene identified to be sequestered by SGs and less translated upon sodium arsenite treatment in human cutaneous melanoma (data from cBioPortal). (**D**) Co-FISH/IF for G3BP1 (IF, red) and *KMT2D* (FISH, cyan) using antisense (top) and sense (bottom) probes with DAPI nuclear counterstain (blue) in A375 cells stressed with 500 µM sodium arsenite for 1 hour. (**E**) Bar chart showing quantification of *KMT2D* FISH puncta colocalization with SGs (n=3). Error bars represent SEM. Significance determined by unpaired T-test. (**F**) IF for G3BP1 (green) and KMT2D (red) with DAPI nuclear counterstain (blue) in A375 cells stressed with 25 µM sodium arsenite for 24 hours. (**G**) Bar chart showing quantification of KMT2D protein levels from IF images of A375 cells stressed with 25 µM sodium arsenite for 24 hours (n=3). AU=arbitrary units. Error bars represent SEM. Significance determined by unpaired T-test. (**H**) Pie chart showing percentage of all protein-coding genes (top) or SG-sequestered and polysome depleted genes (bottom) that encode SOX10-interacting proteins. Significance determined by Fisher’s exact test. (**I**) Kaplan-Meier curve showing tumor onset rate in *MCR:SOX10 & gRNA:NT+tyr* and *MCR:SOX10 & gRNA:kmt2d+tyr* zebrafish. Significance determined by log-rank test. (**J**) IGV tracks showing CUT&RUN peaks for H3K4me1, H3K27ac, and SOX10 at candidate neural crest genes in *MCR:SOX10 & gRNA:NT+tyr* (S) and *MCR:SOX10 & gRNA:kmt2d+tyr* (kS) melanomas. Highlighted regions show increased SOX10 peak intensity in kS melanomas. (**K**) GSEA plot for expression of neural crest cell differentiation genes in *MCR:SOX10 & gRNA:NT+tyr* vs *MCR:SOX10 & gRNA:kmt2d+tyr* melanomas (n=3). Scale bars, 10µm (**D**, **F**).

One gene whose mRNA was sequestered by G3BP1 and less translated upon SG induction was *KMT2D*, a known melanoma tumor suppressor whose protein has been shown to be downregulated during human melanomagenesis (Supp. Fig. 11G)^35^. Loss-of-function mutations in *KMT2D* during development are linked to the neurocristopathy Kabuki Syndrome, where cells fail to differentiate out of the neural crest state^36,37^. This positions *KMT2D* sequestration by SGs as a strong candidate to regulate the neural crest fate adoption of melanoma initiating cells. We performed FISH/IF to confirm *KMT2D* mRNA colocalizes with SGs in human and zebrafish melanoma cells (Fig. 3D-E; Supp. Fig. 12A). We then used RBPscan, an *in vivo* RNA editing tool that reports RNA binding protein motif preference, to quantify the number of G3BP1 binding motifs on *KMT2D* mRNA (Supp. Fig. 12B-C)^38,39^. RBPscan revealed a significant, positive correlation between G3BP1 binding sites and RIP-seq log_2_FC that leveled off at 5+ sites (Supp. Fig. 12B). We found that human *KMT2D* has 17 G3BP1 binding sites, ranking it in the top 0.3% of all expressed mRNAs (Supp. Fig. 12C-D). Zebrafish *kmt2d* contains five G3BP1 binding sites, maintaining its position in the category of mRNAs most vulnerable to SG-mediated sequestration (Supp. Fig. 12B-C). We reasoned that prolonged capture of *KMT2D* mRNA in SGs could alter its protein levels. To test this, we used a chronic, 24-hour sodium arsenite treatment in A375s and found ∼40% decrease in KMT2D protein without alteration to its mRNA levels (Fig. 3F-G, Supp. Fig. 13). We confirmed that dsRNA-induced SGs similarly sequester *KMT2D* mRNA (Supp. Fig. 14). These data indicate that *KMT2D*, a major neural crest-regulating gene, encodes mRNA that is highly vulnerable to G3BP1-mediated sequestration, and this capture is sufficient to reduce its protein levels.

We next sought to understand how SG-mediated KMT2D downregulation could affect neural crest fate acquisition, which is primarily driven by SOX10^3,40,41^. A recent report probed the human melanoma SOX10 interactome^42^. Interrogation of this dataset revealed that *KMT2D*, among seven other SG-sequestered mRNAs (*ATN1, EP400, FOXK1, GLTSCR1, NCOA6, RERE, MED15*), encode proteins found in the top 100 strongest SOX10 interactors^42^ (Fig. 3H). Furthermore, *KMT2D* missense mutations significantly co-occur with *SOX10* alterations in human melanoma, suggesting negative regulation (Supp. Fig. 15A). This led us to hypothesize that KMT2D negatively regulates the ability of SOX10 to activate the neural crest program required for melanoma initiation. We modeled human melanoma genetics in our zebrafish by mosaically knocking out *kmt2d* using our mitfa:Cas9 2xU6:gRNA plasmid in combination with overexpression of human *SOX10* via MCR^43^. We found that SOX10 activity, as read out by abundance of *crestin+* patches per fish^3^, was increased in the *kmt2d*-edited background (Supp. Fig. 15B-C). In agreement, *MCR:SOX10; gRNA:kmt2d+tyr* zebrafish showed faster melanoma onset compared to *MCR:SOX10; gRNA:NT+tyr* controls (Fig. 3I; Supp. Fig. 15D-E). Supporting these observations, CUT&RUN for SOX10 in *kmt2d*-edited melanomas showed increased SOX10 binding at key melanoma and neural crest-regulating genes, including *crestin*, *dlx3b*, and *foxd3* (Fig. 3J). In agreement, RNAseq revealed an upregulation of genes involved in neural crest cell differentiation by GSEA, primarily driven by genes including *dlx3b, ednrba, zeb2b, cdh2,* and *sox10* itself (Fig. 3K). These data establish a model where SGs capture *kmt2d* mRNA to reduce its protein levels, thereby enhancing SOX10 binding at genes crucial for neural crest reprogramming, ultimately driving the acquisition of oncogenic fate.

### Blocking stress granule formation halts melanoma onset and reduces tumor burden

We next asked whether melanomas could develop endogenously without *g3bp1* and its paralog *g3bp2*, which have been shown to diminish SG formation when disrupted in combination^44^. We utilized our mitfa:Cas9 2xU6:gRNA plasmid and injected zebrafish with gRNAs targeting *g3bp1* and *g3bp2*. Tumors expressing gRNAs against *g3bp1/2* showed significantly delayed and reduced CPZ and tumor onset relative to non-targeting gRNA controls (Fig. 4A-B; Supp. Fig. 16A-B). When we profiled the editing of *g3bp1* and *g3bp2* in tumors that did form, we found that only 2/10 tumors were edited for both *g3bp1* and *g3bp2*, whereas 8/10 tumors showed editing for either *g3bp1* or *g3bp2* (Supp. Fig. 16C-F). Integration of both plasmids was detected in 7/8 of the edited tumors, suggesting active selection for clones of tumor cells that retain one functional g3bp (Supp. Fig. 16F-H). We reasoned that delayed melanoma onset may be due to epigenetic dysregulation, especially due to the predicted increase in kmt2d function. Given there are currently no zebrafish-compatible kmt2d antibodies, we performed CUT&RUN for its primary epigenetic mark, H3K4me1^45^. We found a massive increase in H3K4me1 peaks in *g3bp1/2*-edited tumor cells, which is consistent with the predicted increase in kmt2d protein (Fig. 4C). Motif enrichment revealed that these peaks were deposited at stress response genes, including NFE2L2, ATF3, and AP-1 targets, such as *jun* and *chac1* (Fig. 4D; Supp. Fig. 17)^46–48^. Decreased peak abundance was found to be enriched at motifs targeted by core neural crest and melanocyte factors TCF3, PAX3, and MITF, resulting in decreased signal at melanoma identity genes like *tryp1b* and *ednrba* (Fig. 4D; Supp. Fig. 17)^49–51^. Together, these data suggest that g3bp1/2-mediated SG formation is imperative to inform melanoma cell identity for sustained tumorigenesis.

**Figure 4.**
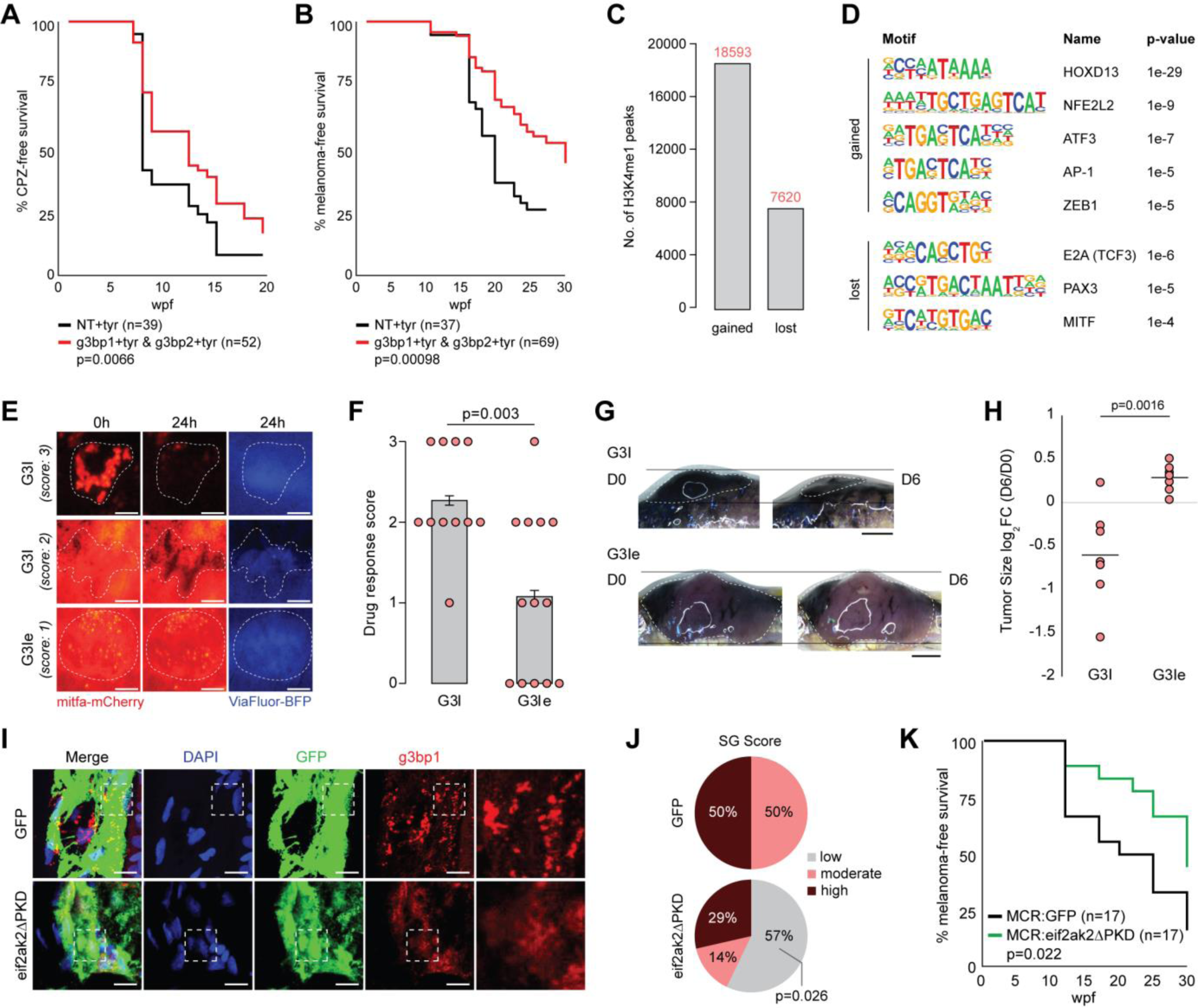
Genetic and chemical disruption of stress granule formation reduces melanoma incidence and tumor burden. (**A-B**) Kaplan-Meier curve showing CPZ (**A**) and tumor (**B**) onset rate in control (*gRNA:NT+tyr*) and *g3bp1/2-KO* (*gRNA:g3bp1+tyr & gRNA:g3bp2+tyr*) zebrafish. Significance determined by log-rank test. (**C**) Bar chart showing number of H3K4me1 CUT&RUN peaks gained and lost in *g3bp1/2-KO* melanoma. (**D**) Motifs enriched at gained (top) and lost (bottom) H3K4me1 peaks in *g3bp1/2-KO* melanoma. (**E**) Representative fluorescent live imaging of zebrafish CPZs injected with G3I or G3Ie at time 0h and 24h with ViaFluor -BFP injection tracer. (**F**) Bar chart showing drug response score of CPZs injected with G3I (n=11) or G3Ie (n=13). 0=no response, 1=partial response, 2=moderate response, 3=complete response. Each dot is one CPZ. Error bars represent SEM. Significance determined by unpaired T-test. (**G**) Representative live images of zebrafish melanomas at day (D) 0 and D6 post-injection with G3I (top) or G3Ie (bottom). (**H**) Quantification of melanoma size at D6 post-injection with G3I (n=7) or G3Ie (n=7) normalized to D0 size. Line represents mean. Significance determined by unpaired T-test. (**I**) IF for GFP (green) and g3bp1 (red) with DAPI nuclear counterstain (blue) in representative *MCR:GFP* (top) and *MCR:eif2ak2ΔPKD-T2A-GFP* (bottom) CPZ. (**J**) Pie chart showing qualitative SG abundance detected in CPZs from *MCR:GFP* (top) (n=8) and *MCR:eif2ak2ΔPKD-T2A-GFP* (bottom) (n=7) zebrafish. Significance determined by Fisher’s exact test. (**K**) Kaplan-Meier curve showing tumor onset rate in *MCR:GFP* and *MCR:eif2ak2ΔPKD-T2A-GFP* zebrafish. Significance determined by log-rank test. Scale bars, 0.1 cm (**E**), 0.25 cm (**G**), 10 µm (**I**).

A small molecule G3BP1/2 inhibitor called G3Ib (here, G3I) was recently generated and provided the opportunity to test whether SGs are a therapeutic vulnerability of melanoma cells^52^. G3I can both dissolve pre-existing SGs and can prevent SG induction (Supp. Fig. 18A-D)^52^. We first tested the function of this compound in human A375 cells by treating them with sodium arsenite alone or in combination with G3I or its inactive enantiomer control (G3Ib’, here G3Ie)^52^. A375s with inhibited SG function showed ∼3.6x decreased viability when compared with cells capable of forming SGs, reinforcing their cytoprotective function^53^ (Supp. Fig. 18E). G3I targets a G3BP1 domain that is evolutionarily conserved between human and zebrafish and was found to be functional in cultured zebrafish melanoma cells (Supp. Fig. 18F-G)^54^. These observations motivated us to inject G3I directly into zebrafish CPZs and melanomas. We found robust and sometimes complete elimination of CPZs injected with G3I, whereas G3Ie injected control CPZs displayed only partial loss of mitfa-mCherry signal (Fig. 4E-F). In agreement, intratumoral delivery of G3I resulted in a significant decrease in tumor size over a week-long administration period, whereas all G3Ie-injected tumors maintained size or grew (Fig. 4G-H). G3I-treated tumors displayed reduced SG staining upon IF for g3bp1, demonstrating that tumor shrinkage was coincident with SG dissolution (Supp. Fig. 19). These findings reveal that melanoma cells leverage SGs for their continued oncogenic competence, and disruption of these condensates is sufficient to eradicate precursor lesions and reduce tumor burden.

Finally, we tested whether disrupting the inducer of SGs, rather than SG components directly, could recapitulate our previous findings. We hypothesized that impairing the molecular cascade triggered by eif2ak2-mediated dsRNA sensing would halt melanoma formation. To test this, we used MCR to overexpress zebrafish *eif2ak2* lacking its protein kinase domain (*eif2ak2ΔPKD*), which allows for dsRNA binding but suppresses autophosphorylation in a dominant negative manner^55,56^. Immunofluorescence for g3bp1 showed diminished SG formation in 4/7 *MCR:eif2ak2ΔPKD-T2A-GFP* CPZs (Fig. 4I-J; Supp. Fig. 20). In turn, *MCR:eif2ak2ΔPKD-T2A-GFP*-expressing zebrafish showed a significant delay in melanoma onset (Fig. 4K). These findings confirm that dsRNA-sensing is a primary SG inducer in CPZs, and activation of this pathway is a central driver of melanoma initiation. Targeting SG assembly directly or blocking inducers of SGs, such as ERV-derived dsRNA, is sufficient to halt tumorigenesis.

## Discussion

Here we use a primary model of zebrafish melanoma to parse the molecular events governing acquisition of oncogenic fate. We report that ERV expression can be used to define stages of melanoma initiation in both zebrafish and human. While it remains untested whether there are distinct biological effects caused by expression of different ERV family-derived elements, we demonstrate that a conserved outcome of their expression, shown as p-eif2ak2-mediated induction of SGs, is a primary tumorigenic effector. To the best of our knowledge, this report is the first demonstration that SGs form endogenously, *in vivo*, upon natural tumorigenic stress in cancer precursor cells and persist in established primary tumors. Supporting SG induction is robust transcriptional upregulation for key SG regulators in both zebrafish and human. This transcriptional control may serve to increase the abundance of SG components to lower the threshold in which stress induces their formation, since relative stoichiometric alterations can induce SG formation independent of cell stress, or to maintain their persistence^44,57–59^. Enhancing this process by overexpressing *hG3BP1 in vivo* demonstrates an evolutionarily conserved, pro-tumorigenic role for upregulating the expression of core SG genes.

We sought to mechanistically define how SG induction could impart oncogenic reprogramming. The predicted outcome of mRNA sequestration and decreased translation is perturbation to the pathways in which their proteins participate. We report the main biological pathway enriched by SG-controlled mRNAs in melanoma is epigenetic regulation directly implicating putative SOX10 interactors. *KMT2D* emerged as a primary candidate whose suppression by SGs could impart melanoma fate. In Kabuki Syndrome and cancer, most *KMT2D* alterations are loss-of-function mutations^60,61^. However, complete knockout of *KMT2D* is embryonically lethal in both mouse and zebrafish^62,63^. We suspect there is a baseline expression level of KMT2D required to maintain cell viability, but a level at which depletion becomes tumorigenic. Capturing of *KMT2D* mRNA by SGs reduced its protein levels by ∼40%, which matched the range of indels seen in *kmt2d* edited tumors. As a direct SOX10 interactor, KMT2D likely acts to restrict SOX10 from activating the neural crest program, and its loss leads to persistence of the SOX10-driven neural crest state in Kabuki Syndrome and acquisition of neural crest-like state in melanoma^42^. It is a compelling hypothesis that similar chromatin remodeling imparted by SG-mediated mRNA regulation could facilitate lineage plasticity in other tumors by relieving their respective embryonic lineage factors. More broadly, our SG-regulated set includes other major controllers of pluripotency regulation, such as *DOT1L*, whose knockdown enhances stem cell reprogramming^64,65^. SGs likely facilitate acquisition of cancer stemness *in vivo* through multiple pathways.

Our genetic data demonstrates that there is abundant tumorigenic stress and a requirement for full SG competence to generate melanoma, as evidenced by selection for tumor clones that retain a functional *g3bp* allele. We have previously observed this phenomenon for other essential melanoma genes including *sox10*^3^. Edited tumors showed chromatin reprogramming at sites controlled by master stress-response regulators like NFE2L2, ATF3, and AP-1, and melanoma resistance programs controlled by ZEB1^46–48,66^. The concomitant loss of H3K4me1 at melanocyte and neural crest-defining motifs reveals a fundamental rewiring of cell identity, underscoring SGs as both mediators of stress tolerance and fate controllers. Despite these findings, using genetic editing to disrupt SG assembly is clinically impractical. We therefore leveraged the G3BP1/2 inhibitor G3I to determine whether small molecule-mediated SG dissolution could serve as a therapeutic strategy *in vivo*^52^. G3I eradicated CPZs and reduced tumor burden, establishing SG inhibition as a therapeutically viable approach. Melanoma is the deadliest form of skin cancer in part due to late-stage diagnosis, underscoring the importance of early intervention^67^. The presence of SGs in human nevi combined with the ability of G3I to abolish CPZs positions SG inhibition as a preventative melanoma therapy. This application extends beyond melanoma, as SGs have been implicated in tumor initiation and therapy resistance in multiple solid tumor types, including pancreatic ductal adenocarcinoma, hepatocellular carcinoma, sarcoma, and lung cancer^68–72^. Combining our findings with these observations provokes significant enthusiasm for the use of SG-inhibiting molecules to exploit a pan-cancer cell property required for tumor formation and persistence. Further contributing to the goal of cancer prevention, our identification of stage-defining ERV signatures offers potential rapid diagnostic utility to distinguish lesions that may be prone to malignant transformation. The ERV-SG axis presents a promising avenue to diagnose and treat cancer.

## Methods

### Zebrafish husbandry

The following zebrafish lines were used in this study: wild-type *TU*, wild-type *casper*; *mitfa:hBRAF^V600E^*, *mitfa^-/-^*, *p53^-/-^* (triples); *mitfa:hBRAF^V600E^*, *mitfa^-/-^*, *p53^-/-^*, *crestin:eGFP* (tCE). Animals were maintained at Boston Children’s Hospital. All experiments and husbandry were performed in accordance with protocols approved from Boston Children’s Hospital’s Institutional Animal Care and Use Committee (IACUC).

### Zebrafish embryo injections

For melanocyte-specific overexpression models, single cell-stage embryos were injected with equivalent concentrations (range 20-40 ng/µL) of the described MCR plasmids in addition to 20 ng/µL mitfa:mCherry, 10 ng/µL alb gRNA, 0.2 mg/mL Cas9 protein, and 25 ng/µL Tol2 mRNA. For mosaic, melanocyte-specific knockout models, single cell-stage embryos were injected with equivalent concentrations (final 40 ng/µL) of mitfa:Cas9 2xU6:gRNA plasmids, 20 ng/µL mitfa:mCherry, and 25 ng/µL Tol2 mRNA. Control plasmids contained an empty first site with a *tyr* gRNA in the second site, and experimental plasmids contained a gRNA against the targeted gene in the first site and *tyr* gRNA in the second site, where tyr loss eliminates pigmentation to facilitate imaging. The gRNA sequences used are as follows: *alb* 5’ – GGTTTGGGAACCGGTCTGAT – 3’; *tyr* 5’ – GGACTGGAGGACTTCTGGGG – 3’; *kmt2d* (gRNA1) 5’ – GAGGAGCCATCACCCGACTG – 3’; *kmt2d* (gRNA2) 5’ – GTATTGACTGTGGCATGCGA – 3’; *g3bp1* 5’ – GACCGGATAGGAGGGTCAAGG – 3’; *g3bp2* 5’ – GGATTCGCTGGTCACGTGGA – 3’. Embryos were incubated at 28°C in E3 and screened for melanocyte rescue via mCherry signal at 5 dpf using a Nikon SMZ18 microscope.

### Zebrafish imaging

Zebrafish were anesthetized with 4% tricaine (MS-222) and imaged on a 1.8% E3-agarose mold using a Nikon SMZ18 microscope. For imaging Ubi:hG3BP1-mCherry, larvae were mounted in 0.8% low-melting point agarose containing 0.16 mg/mL Tricaine in 6-well, glass-bottom plates. Wells were filled with E3 containing 0.16 mg/mL Tricaine. The plate was placed in an Oko Lab Microscope Incubator set to 42°C for one hour with acquisition of brightfield and mCherry every 10 minutes. Images were minimally processed with FIJI and Adobe Photoshop.

### Cloning

To create MCR:G3BP1-GFP and MCR:eif2ak2ΔPKD-T2A-GFP, canonical human G3BP1 CDS (Ensembl ENST00000356245.8) and zebrafish eif2ak2 CDS lacking protein kinase domain (Ensembl ENSDART00000164338.2; protein domains determined using NCBI BlastP) were ordered as gBlocks from IDT and cloned into middle entry vectors using pENTR/D-TOPO Cloning Kit (ThermoFisher cat. K240020) per manufacturer’s protocol. Final MCR constructs were generated using Gateway LR Clonase II Enzyme Mix (ThermoFisher cat. 11791020) per manufacturer’s recommendations. Plasmid sequences were confirmed using Nanopore Rapid Plasmid Sequencing services provided by Plasmidsaurus and Quintara.

Mitfa:Cas9 2xU6:gRNA constructs were generated by digesting the base plasmid (Addgene cat. 118844) with BseRI (NEB cat. R0581) for cloning gRNA into site 1 and AarI (ThermoFisher cat. ER1581) for cloning gRNA into site 2. gRNAs were designed by CHOPCHOP. gRNA efficiency was tested first by cloning into Ubi:Cas9 U6:gRNA using BseRI^73^. Wild-type *TU* embryos were injected and collected at 48 hpf for gDNA isolation. PCR was performed using primers (Supp. Table 1) flanking the targeted cut-site and products were purified via Qiagen’s QIAquick PCR Purification Kit (cat. 28106) and sequenced by Psomagen. CRISPR-editing efficiency was calculated using Synthego’s ICE web tool. gRNAs with >10% editing in embryos were used for adult melanoma generation and analysis. Editing was secondarily validated using T7 endonuclease I (T7E1) assay, where PCR products were re-annealed and digested with T7E1 (NEB, cat. M0302L) for one hour at 37°C, followed by standard agarose gel electrophoresis.

### Cell culture

Human A375 cells (ATCC cat. CRL-1619) were incubated at 37°C and 5% CO_2_ in DMEM with 10% FBS, 1% Pen/Strep (Corning cat. 30-002-CI), and 1% L-Glutamine (ThermoFisher cat. 25030081). Zebrafish melanoma cells^54^ were incubated at 28°C and 5% CO_2_ in DMEM with 10% FBS, 1% Pen/Strep/GlutaMAX (ThermoFisher, cat. A5873601), and 1% L-Glutamine. All cells were passaged readily and maintained with minimal total passage number (<10).

### Histology

Zebrafish were euthanized and placed in 10% formalin for 48 hours at 4°C. Paraffin embedding and sectioning were performed by Brigham & Women’s Histopathology Core. Immunohistochemistry was performed using Proteintech IHC Prep & Detect Kit for Rabbit Primary Antibody (cat. PK10017) per manufacturer’s recommendations. Briefly, 5 µm sections were deparaffinized with xylene and rehydrated using an ethanol series. Sections were incubated in Antigen Retrieval Buffer at 98°C for 20 minutes. Endogenous peroxidase activity was suppressed using Quenching Buffer. Sections were incubated with primary antibodies (Rb α mCherry, Abcam ab167453; Rb α p-ERK, CST 4370; Rb α eif2a, CST 5324S; Rb α p-eif2a, CST 9721S; Rb α eif2ak2, Abcam ab184257; Rb α p-eif2ak2, Abcam ab32036) at 1:100 overnight at 4°C. Primary antibody was thoroughly rinsed, after which sections were incubated with HRP-Goat anti-Rabbit Secondary Antibody for 30 minutes at RT. Chromogenic signal was developed for 5-10 minutes (stopped relative to onset of background signal). Sections were counterstained with hematoxylin Counter Staining Reagent and then dehydrated in an ethanol series. Sections were briefly immersed in xylene then mounted with Mounting Media. Images were acquired using a Nikon Eclipse E600 and were minimally processed using FIJI.

For zebrafish section immunofluorescence, 5 µm sections were deparaffinized with xylene and rehydrated using an ethanol series. Sections were incubated in Citric Acid Based Antigen Unmasking Solution (Vector Laboratories; cat. H-3300) for 30 minutes at 98°C. After, sections were blocked using 0.2% BSA and incubated with primary antibodies (Ch α mCherry, Novus NBP2-25158; Rb α g3bp1, Proteintech 13057-2-AP; Rb α stau2, Proteintech 15998-1-AP; Rb α tia1l, Novus NBP1-79932; Gt α GFP, Abcam ab5450) at 1:300 overnight at 4°C. Primary antibody solution was washed thoroughly and then sections were incubated in secondary antibodies (G α Ch 488, Invitrogen A11039; Gt α Rb 568, Invitrogen A11011; D α Gt 488, Invitrogen A11055; D α Rb 568, Invitrogen A11008) at 1:500 for 3 hours at RT. Sections were counterstained with DAPI and mounted with ProLong Diamond Antifade Mountant (Invitrogen; cat. P36961). Human nevi were processed using the same protocol with the following primary antibodies (Rb α SOX10, CST 89356S; Gt α SOX10, R&D af2864; Rb α G3BP1, Proteintech 13057-2-AP; Ms α G3BP1, Abcam ab56574; Ms α J2, CST 76651L). Images were acquired using a Leica TCS SP8 and were minimally processed using FIJI.

For cell culture immunofluorescence, cells were grown in 24-well glass bottom plates, treated as described in text, and then fixed for 10 minutes with 4% PFA. Cells were washed briefly with 0.2% Triton X-100 in PBS and then blocked with 1% BSA for 1 hour at RT. Primary antibodies (Ms α G3BP1, Abcam ab56574; Rb α G3BP1, Proteintech 13057-2-AP; Gt α TIA1, Invitrogen PA5-18699; Rb α KMT2D, Millipore Sigma HPA035977; Rb α EIF4G1, Abnova H00001981-1-A01; Rb α PABP, Abcam ab21060) were then added at 1:500 for 1 hour at RT. Cells were then washed with 0.1% Tween-20 and incubated with secondary antibodies (Gt α Ms 568, Invitrogen A11004; Gt α Rb 488, Invitrogen A11008; D α Gt 488, Invitrogen A11055; D α Rb 568, Invitrogen A10042; D α Ms 647, Invitrogen A31571) at 1:1000 for 1 hour at RT. After, cells were washed with 1xPBS/DAPI and stored at 4°C in 1xPBS. Images were acquired using a Nikon Eclipse Ti Spinning Disk Confocal and were minimally processed with FIJI.

For cell culture immunofluorescence in combination with fluorescent *in situ* hybridization, cells were grown in 6-well plates on glass coverslips. Cells were fixed using 2% PFA for 20 minutes at RT. Cells were dehydrated then rehydrated using a symmetrical 25-50-75-100% methanol/PBS+0.1% Tween-20 series. Cells were incubated with pre-heated hybridization buffer (10mM Tris pH 7.4, 600 mM NaCl, 1 mM EDTA, 0.25% SDS, 5% Dextran Sulfate, 1x Denhardt’s, 200 µg/mL Yeast tRNA, 50% DI Formamide, up to 20mL with DEPC H_2_O) with 1:100 probe for 16 hours at 65°C in an incubation chamber with paper towels soaked in 5X SSC/50% DI Formamide. Coverslips were washed briefly with pre-heated 5X SSC then incubated with 0.2X SSC twice for 20 minutes. After, cells were washed with TN (100 mM Tris pH 7.5, 150 mM NaCl) with 0.05% Tween-20 (TN+T) then blocked with TN with 10% HISS and 0.05% w/v Akoya Blocking Reagent Powder (cat. FP1020) (TN+B) for 30 minutes at RT. Anti-DIG-POD (Millipore Sigma cat. 11207733910) was then added at 1:1000 in TN+B for 1 hour at RT. Cells were washed with TN+T and then incubated in tyramide working solution (Akoya TSA Plus Cyanine 5 cat. NEL745001KT) at 1:100 for 30 minutes at RT. Signal was quenched using 4% PFA for 30 minutes at RT. For co-immunofluorescence, cells were then washed with 0.2% Triton X-100 in PBS, after which the previously detailed protocol was followed. Coverslips were mounted with ProLong Diamond Antifade Mountant (Invitrogen; cat. P36961) on glass slides. Images were acquired using a Nikon Eclipse Ti Spinning Disk Confocal. *In situ* probes were prepared using in vitro transcription of PCR-amplified targets with T7 promoter attached to the forward or reverse primer for downstream sense and antisense transcription, respectively (Supp. Table 1). 300 ng of gel extracted PCR product was incubated with T7 polymerase (Millipore Sigma cat. 10881767001), DIG RNA labeling mix (Millipore Sigma cat. 11277073910), and RNase inhibitor (Millipore Sigma cat. 3335399001) for 2 hours at 37°C. DNA was digested using DNase I (Millipore Sigma cat. 4716728001) for 15 minutes at 37°C. RNA probes were precipitated by mixing 10 µL 4M LiCl, 1 μL Glycogen (2 µg/μL), 100 µL TE-DEPC, and 400 µL 100% ethanol with overnight incubation at -20°C. RNA probes were pelleted, washed with 100% ethanol, dried, and reconstituted in DEPC H_2_O, then stored at -20°C in 10% DEPC H_2_O/90% DI Formamide.

For H&E staining, 5 µm sections were deparaffinized and processed using Vector Laboratories Hematoxylin & Eosin Stain Kit (H-3502), per manufacturer’s protocol. Briefly, sections were rehydrated, stained with Hematoxylin, Blueing Reagent, and Eosin Y, then dehydrated in ethanol and mounted with VectaMount Express Mounting Medium (H-5700). Images were acquired using a Nikon Eclipse E600 and were minimally processed using FIJI.

### Image quantification

SGs per nuclei, percent of cells containing SGs, and cells per field of view analyses were manually quantified using Imaris. Quantification of KMT2D FISH puncta that colocalize with SGs was determined using the FIJI *Image Calculator* tool with subtraction of nuclear signal using DAPI overlap. KMT2D protein intensity was quantified using the FIJI *Measure* tool.

### Zebrafish melanoma RNAseq

For transposable element (TE) element expression of zebrafish melanoma initiation, our previously generated RNAseq^6^ was analyzed using TEtranscripts^74^ with default parameters. Differential TE expression was determined using |log_2_FC|>1 and p-adj.<0.05. SQuIRE^75^ (v0.9.9.92) was also used to quantify TE expression using the same dataset using default settings with differential expression quantified using |log_2_FC|>1 and p-adj.<0.05 via DESeq2 (v1.42.1) with assistance from Claude AI. TE coordinates and family information was obtained via UCSC danRer11 RepeatMasker annotation. MITF binding motif occurrences were identified +/-5kb of *zferv* elements using FIMO with JASPAR MA0620.2 with p<10e-4 cut-off.

For RNAseq analysis of MCR:SOX10; gRNA:NT+tyr (n=3) and MCR:SOX10; gRNA:kmt2d+tyr (n=3) melanoma, tumors were bulk harvested and digested in Trizol. RNA was collected using Direct-zol RNA Miniprep Plus Kit (Zymo, cat. R2052) per manufacturer’s protocol. Library prep and sequencing was performed by Plasmidsaurus, with >12M pA reads obtained per sample. Transcripts were quantified with Salmon (v1.10.3) and DESeq2 (v1.42.1) was used for differential gene expression analysis. GSEA analysis was performed using fgsea (v1.28.0) using gene sets from MSigDB via msigdbr (v25.1.0). Analyses were performed with assistance from Claude AI.

### Human melanoma RNAseq

Raw reads were downloaded from the Gene Expression Omnibus (accession GSE112509)^30^. The cohort is laser-microdissected human nevi (n=23) and primary melanomas (n=57) sequenced as poly-A selected reverse-stranded libraries, 76 bp paired-end, on an Illumina HiSeq 2000. Interleaved FASTQ files were split and trimmed with fastp 0.20.1 (--interleaved_in --trim_poly_g --disable_quality_filtering).

The STAR 2.7.10a index was generated from the NCBI RefSeq GRCh38.p14 assembly (GCF_000001405.40) with --sjdbGTFfile set to the matching RefSeq GTF and --sjdbOverhang at its default of 100. Reads were aligned with STAR 2.7.10a using --outFilterMultimapNmax 100 --winAnchorMultimapNmax 100 --outSAMattributes NH HI AS nM --outSAMtype BAM SortedByCoordinate, all other parameters at defaults. The coordinate-sorted BAM files written by STAR were indexed with samtools.

Gene and transposable element counts were generated together with TEcount (TEtranscripts 2.2.4) using -- mode multi --stranded reverse. The gene annotation was the NCBI RefSeq annotation of GRCh38.p14; gene identifiers are HGNC symbols. The repeat annotation was the RepeatMasker-derived TEtranscripts annotation on the same assembly. The matrix contains 50,037 gene rows and 1,330 repeat rows. Repeats are quantified at subfamily level (subfamily:family:class), aggregating all copies; no locus-level inference is made.

Of 80 published libraries, 74 completed quantification and 73 had matching clinical annotation and were carried forward. Serial sections of the same lesion were summed with collapseReplicates, giving 68 lesions (22 nevi, 46 melanomas) from 53 patients, 15 of whom contributed both a nevus and a melanoma.

featureCounts (subread 2.1.1) was run twice over the same BAM files and the same RefSeq annotation, once with -t exon and once with -t gene, all other flags identical (-g gene_id -p -s 2 -T 16). The exonic rate is exon-assigned reads divided by gene-body-assigned reads per library, and at lesion level the ratio of the two sums across sections.

Where an analysis is reported as adjusted for the exonic rate, it enters the model as a covariate, or is removed from both variables by linear regression before the residuals are correlated. Each test states which.

Size factors were estimated from the gene count matrix and applied to the TE count matrix. Features were retained at 5 or more counts in at least 9 lesions, 9 being the size of the smallest of the four status-by-subtype groups, leaving 1,122 TE subfamilies and 26,932 genes. Differential expression used DESeq2 1.50.2 (R 4.5.2) with the paired design ∼ patient + Status across the 15 patients contributing both lesions, at a Benjamini-Hochberg FDR of 0.05. Significance was assessed across all 1,122 TE subfamilies; of the 27 subfamilies passing FDR 0.05, the 20 belonging to the LTR class were carried forward as the signature (7 higher in melanoma, 13 higher in nevi). The seven significant, non-LTR subfamilies took no part in the score. Variance-stabilized counts (VST, blind) were used for principal component analysis, correlation and visualization. As a sensitivity check the paired log fold changes were recomputed with size factors estimated from the TE matrix itself. Two per-lesion measures were taken forward from this result: i) total LTR abundance, which is the mean variance-stabilized expression across the 565 LTR subfamilies passing the filter, and ii) the composition score, which is the log ratio of the geometric mean of the subfamilies significantly higher in melanoma to the geometric mean of those significantly higher in nevi, computed from raw counts with a pseudocount of 0.5; it was recomputed with pseudocounts of 1 and 5 and with subfamilies carrying any zero count excluded. Within-pair analyses regressed the change in each measure on the change in exonic rate.

The subfamilies entering the ratio were selected using only the 30 lesions of the 15 paired patients. The remaining 38 lesions (7 nevi, 31 melanomas), from patients contributing a single lesion, took no part in selection. A classification threshold was chosen on the 30 selected lesions by maximizing Youden’s J and applied unchanged to the 38; AUC is reported in preference to raw accuracy.

Monte Carlo significance was assessed with empirical P values computed as (k + 1) / (N + 1) under two resamplings, each of 10,000 iterations: nevus/melanoma labels permuted within the held-out set, and ratios built from LTR subfamilies drawn at random from those not entering the ratio, matched in number to each of its two sides.

EIF2AK2 was tested on VST counts as a Wilcoxon rank-sum test across all 68 lesions.

### scRNAseq

Data from our previously generated scRNAseq atlas of zebrafish melanoma initiation was analyzed using Seurat’s AddModuleScore. The genes in the SG module were obtained from Yang et al., 2020^44^. The zebrafish orthologs expressed in the dataset and used for quantification are: *hnrnpa2b1, polr2b, usp10, eif3i, eif3d, eif3eb, ddx3xb, eif3g, ppp2r1ba, upf1, tia1, tial1, ddx19a, nup98, caprin1a, kpnb1, g3bp1, g3bp2, csde1, taf15, trim56, hdac6, ppp1r10, ubap2l, nufip2, atxn2l, atxn2, fmr1, prrc2c, srsf3a, ubap2a, trim25, cbx1a, macf1a*.

### CUT&RUN

For H3K27ac CUT&RUN on CF, CPZ, and melanoma cells, zebrafish were harvested to obtain the tissue of interest. The tissue was digested via rapid chopping for 1-2 minutes in TrypLE Express (ThermoFisher, cat. 12605028), followed by incubation in a Thermomixer set at 37°C, 400 rpm for 45 minutes. Samples were pipette mixed to generate single cell suspensions then filtered with a 100 µm filter. Samples were spun at 500 rcf for 5’ at 4°C and resuspended in FACS buffer (2% FBS, 1% Pen/Strep, 7% ultrapure water, 90% 1x PBS) twice. Samples were then filtered through the lid of FACS tubes (Corning, cat. 352235) and processed using a BD FACSAria to collect mCherry+ (CF or tumor) and mCherry^high^ (CPZ) cells with assistance from Boston Children’s Hospital’s Flow Cytometry Core. Approximately 5000 CF and 2000 CPZ cells were pooled from N=6 zebrafish skin, and 100000 tumor cells were collected from pooled N=3 tumors. All collected cells were used for input and evenly divided among reactions. For H3K4me1, H3K27ac, and SOX10 CUT&RUN of MCR:SOX10; gRNA:NT+tyr (n=2) and MCR:SOX10; gRNA:kmt2d+tyr (n=2) melanomas, tumors were bulk harvested and processed for FACS without undergoing sorting, as hSOX10 is only overexpressed in melanocytes. 100000 cells were used per reaction. For H3K4me1 CUT&RUN of gRNA:NT+tyr (n=2) versus gRNA:g3bp1+tyr, gRNA:g3bp2+tyr melanomas (n=1), tumors were bulk harvested and processed for FACS without sorting, with 10% of cells used for genotyping and remainder split equally for reactions. CUT&RUN was performed using CUTANA CUT&RUN Kit v6 (cat. 14-1048) per manufacturer’s recommendations. Briefly, cells were pelleted, resuspended in Wash Buffer, then adsorbed to activated ConA beads. Cell-bead slurries were incubated with 0.5 µg primary antibody (Rb α H3K4me1, Epicypher cat. 13-0057; Rb α H3K27ac, Epicypher cat. 13-0059; Rb α SOX10, CST cat. 89356S; Rb α IgG, Epicypher cat. 13-0042) in Antibody Buffer overnight at 4°C on a nutator.

After, cells were washed in Cell Permeabilization Buffer, incubated with pAG-MNase for 10 minutes at RT, then incubated with 100 mM CaCl_2_ for 2 hours at 4°C. Reactions were neutralized with Stop Buffer at 37°C for 10 minutes. Supernatants containing CUT&RUN-enriched chromatin fragments were then transferred to new tubes. CUTANA DNA Purification Beads were used to purify the DNA, which was eluted in 0.1x TE. 5 ng (or total, if less) CUT&RUN DNA was used as input for CUTANA CUT&RUN Library Prep (cat. 14-1001), which was carried out using manufacturer’s guidelines. Briefly, DNA was end-repaired followed by adapter ligation and U-excision. Modified DNA was purified using SPRIselect bead selection and eluted in 0.1x TE. Samples were indexed with unique i7/i5 primer combinations and PCR amplified using 14 cycles. Resulting libraries were purified using SPRIselect bead selection and eluted in 0.1x TE. Library size and integrity was evaluated using Agilent Tapestation with D1000 ScreenTape (cat. 5067-5582), per manufacturer’s recommendations, to confirm peak enrichment from 200-700 bp. Samples were then pooled at equimolar ratios and sequenced at 2x150 bp paired-end reads with 360 million total read depth via Quintara Biosciences.

CUT&RUN sequences were aligned to the zebrafish genome (Ensembl build version GRCz11) using Bowtie2 v2.2.1 with the following parameters: --local, -N0, -L20. MACS2 v2.1.0 was used to call peaks with the following parameters: --nomodel, --shift -75, --extsize 150, and a q-value threshold of enrichment of 0.01 for all datasets. Motif analysis was performed using HOMER findMotifsGenome.pl with parameters -size 200 -mask, with assistance from Claude AI.

### CPZ & tumor tracking

CPZ and tumor Kaplan-Meier curves were generated in R using the *ggsurvplot* function from the *survminer* package, which calculates significance using a log-rank test. Expression of constructs was validated through phenotypic confirmation for overexpression constructs (ex: GFP fluorescence detected in MCR:GFP and MCR:hG3BP1-GFP CPZs and tumors) or genotyping for mitfa:Cas9 2xU6:gRNA constructs. Primers used for genotyping can be found in Supp. Table 1. PCR products were purified using Qiagen’s QIAquick PCR Purification Kit (cat. 28106) and Sanger sequenced by Psomagen. CRISPR-editing efficiency was calculated using Synthego’s ICE Analysis web tool.

### RIP-seq

A375s were grown to 90% confluency in 15 cm dishes (one per replicate) and treated with 500 µM sodium arsenite or equal volume vehicle (water) in pre-warmed cell culture media for one hour. Cells were washed once with warm media and then once with ice-cold 1x PBS. Cells were scraped in 5 mL ice-cold 1x PBS and pelleted at 1500 rpm for 5 minutes at 4°C. RIP was then performed using the MAGNA RIP RNA-Binding Protein Immunoprecipitation Kit (Millipore Sigma cat. 17-700) per manufacturer’s recommendations. Briefly, magnetic beads were washed in RIP Wash Buffer and incubated with 4 µg antibody (Rb α G3BP1, Proteintech 13057-2-AP; Rb α IgG, Proteintech 30000-0-AP) on a nutator for 30 minutes at RT. Beads were washed and resuspended in 900 µL RIP IP Buffer. Pelleted cells were resuspended in 250 µL RIP Lysis Buffer and centrifuged at 14000 rpm for 10 minutes at 4°C. 100 µL of supernatant was added to the bead-antibody slurry; 10 µL was kept as input RNA; and 10 µL was kept as input for Western blot validation. Reactions were incubated at 4°C overnight on a nutator. Samples were then washed thoroughly with ice-cold RIP Wash Buffer and resuspended at a volume of 500 µL, of which 50 µL was removed for Western blot validation of the IP. Reactions and input RNA samples were then incubated in a Thermomixer at 55°C for 30’ in Proteinase K Buffer. After, samples were briefly spun, placed on a magnet, and supernatants were moved to new tubes. RIP Wash Buffer was added to each sample to bring the final volume to 400 µL, to which 400 µL phenol:chloroform:isoamyl alcohol (125:24:1) was added. Samples were thoroughly vortexed and spun at 14000 rpm for 10 minutes at RT. The aqueous phase was moved to a new tube and mixed with 400 µL chloroform. Samples were spun again at 14000 rpm for 10’ at RT. The aqueous phase was removed, placed in a new tube, and incubated with MAGNA salt solutions, precipitate enhancer, and 850 µL 100% ethanol overnight at -80°C. Samples were spun at 14000 rpm for 30’ at 4°C, and RNA pellets were washed with 80% ethanol. Pellets were spun once more at 14000 rpm for 15’ at 4°C, dried, and resuspended in 10 µL DEPC-H_2_O. RNA integrity and concentrations were determined using Agilent Tapestation with High Sensitivity RNA ScreenTape (cat. 5067-5579). Ribo-depletion, library preparation, and sequencing was performed by Azenta GENEWIZ. Quality control of RNA-seq datasets was performed using FastQC and CutAdapt to remove adaptor sequences and low-quality regions. High-quality reads were aligned to GRCh38 using STAR 2.7.0. Raw read counts were calculated by HTSeq. Differential expression analysis was performed using DEseq2.

### Western blot

For RIP-seq IP validation, input and IP samples were resuspended in Pierce RIPA Buffer (ThermoFisher cat. 89901) with 1x Roche cOmplete Mini, EDTA-free Protease Inhibitor (Millipore Sigma cat. 11836170001). Samples were run using Novex 4-20% Tris-Glycine Plus WedgeWell Gels (Invitrogen cat. XP04202BOX) and transferred using Invitrogen iBlot2 (cat. IB21001) with iBlot2 PVDF Mini Stacks (cat. IB24002). Blots were washed in 1x TBS with 1% Tween-20 (TBST) and blocked using 5% milk in TBST for 1 hour at RT. Primary antibodies (Ms α G3BP1, Abcam ab56574; Ms α GAPDH, Proteintech 60004-1-Ig) were added at 1:1000 overnight at 4°C with gentle rocking. Blots were then washed thoroughly and incubated with secondary antibody (Anti-Ms IgG HRP-linked, CST cat. 7076S) at 1:1000 for 1 hour at RT with gentle rocking. After, blots were washed thoroughly and then incubated for 1 minute with Pierce ECL reagents (ThermoFisher cat. 32209) and developed on autoradiography film (Genesee Scientific cat. 30-101L). Blots were minimally processed using FIJI.

For zebrafish skin and tumor protein expression analysis, tissue was extracted and forcefully chopped for 1-2 minutes with RIPA buffer with 1x protease inhibitor. Samples were then sonicated using Diagenode Bioruptor Next Gen Millenium 3 with 6 cycles at 30 seconds on, 30 seconds off. Samples were centrifuged at 14000 rpm for 10 minutes at 4°C and supernatants were moved to new tubes. Protein concentration was determined using Pierce BCA Protein Assay Kit (ThermoFisher cat. 23227). 20 µg protein was analyzed by Western blot as described above using primary antibodies (Rb α eif2ak2, Abcam ab184257; Rb α p-eif2ak2, Abcam ab32036; Ms α b-Actin, Proteintech 66009-1-Ig) at 1:1000 and secondary antibody (Anti-Ms IgG HRP-linked, CST cat. 7076S; Anti-Rb IgG HRP-linked, CST cat. 7074S) at 1:1000. Blots were minimally processed and protein expression was quantified using FIJI.

### Polysome-seq

A375 cells were grown to 90% confluency in 15 cm dishes (one per replicate) and treated with 500 µM sodium arsenite or equal volume vehicle (water) in pre-warmed cell culture media for one hour. Cells were washed once with warm media and then incubated in media with 100 µg/mL cycloheximide (ThermoFisher cat. J66004.XF) for 5 minutes at 37°C. Cells were washed twice with ice-cold PBS containing 100 µg/mL cycloheximide and then scraped in 5 mL ice-cold PBS and collected in 50 mL conicals. Cells were pelleted by centrifuging at 300 x g for 5 minutes at 4°C. Cells were resuspended in 425 µL hypotonic buffer (50mM Tris HCl pH 7.5, 2.5 mM MgCl _2_, 1.5 mM KCl, 1x protease inhibitor, up to 1mL with 1x PBS) then spiked with 5 µL 10 mg/mL cycloheximide, 1 µL 1M DTT, and 2 µL RNase inhibitor and vortexed. After, 25 µL 10% Triton X-100 and 25 µL 10% sodium deoxycholate were added to each sample, which were then vortexed and centrifuged at 20000 x g for 20 minutes at 4°C. A260 was quantified and 20 O/D units were moved per sample to new tubes that were brought to 500 µL total volume with hypotonic buffer. Polysome profiling was then performed as per Ietswaart et al., 2024^76^. Briefly, cell lysates were layered over 10-50% sucrose gradients (20 mM HEPES pH 7.6, 100 mM KCl, 5 mM MgCl_2_, 10 µg/mL cycloheximide, 1X protease inhibitor cocktail, RNase inhibitor) and centrifuged for 2 hours at 35000 rpm in an SW41Ti rotor (Beckman). Gradients were separated into 12 fractions using a Gilson FC203B fraction collector, with RNA absorbance monitored using a BioComp153 gradient station ip (BioComp Instruments) using an FC-2 Triax flow cell and associated software (v1.53A, BioComp). Fractions 8-12 were pooled as the polysome fractions, ethanol precipitated, and extracted with Trizol following the manufacturer’s directions. In parallel, RNA was Trizol extracted from 5% of the input material. RNA was DNase-treated (TURBO DNase, Invitrogen) and cDNA libraries were constructed from 1 µg input and polysome RNA using the NEBNext Ultra II Directional RNA Library Kit (cat. E7760). Libraries were sequenced using 100 bp paired-end reads on a NextSeq 500 via Tufts genomics core facility.

### In vivo drug treatment

G3I or G3Ie was mixed 4:1 with Biotium ViaFluor SE 405 (cat. 30068). 1 µL of drug mix was injected subcutaneously or intratumorally using a 26g Hamilton Syringe. The syringe was cleaned with 70% ethanol and ultrapure water between each injection. CPZ response was manually scored double-blind. Tumor size was quantified by tracing melanomas and quantifying traced area in Adobe Illustrator using the *measure* tool.

### Tumor transplants

*Casper* adult zebrafish that were between six and 11 months post fertilization were used as transplant recipients. These adult fish received split-dose irradiation of 14 Gy each for two consecutive days prior to transplant. Control and MCR:G3BP1 tumors from adult zebrafish were dissociated as described above and counted with a hemocytometer. Cells were centrifuged at 500 x g for 5 minutes at 4°C to pellet the cells. Cells were resuspended through serial dilution in phenol red-free Matrigel (Corning catalog # 356237) to concentrations of 10000, 5000, 3000, 1000, or 500 cells/µL. Each recipient fish received bilateral subcutaneous injections (1 µL of cells in Matrigel per injection) with cells from control tumors injected on the left side and cells from MCR:G3BP1 tumors injected on the right side. Fish were imaged as described above and limit dilution calculations and curves were generated using the Extreme Limiting Dilution Analysis software (http://bioinf.wehi.edu.au/software/elda/) on data collected 28 days post-transplant.

### RBPscan

Human transcript sequences and CDS coordinates were obtained from NCBI for all protein-coding genes detected in the G3BP1 RIP-seq dataset. For genes with multiple transcripts, the longest transcript per gene was retained. The top 7 G3BP1 binding motifs identified by RBPscan were searched against each transcript sequence. Total binding sites per transcript were summed across all motifs and categorized into bins (0, 1, 2, 3, 4, 5–10, >10 sites). RIP-seq log_2_FC values were assigned per gene and empirical cumulative distribution functions were plotted. Statistical significance was assessed by Kolmogorov-Smirnov test comparing each site-count group against transcripts with no predicted binding sites, with p-values adjusted using the Benjamini-Hochberg correction. Full-length transcript sequences and CDS coordinates for human (ENST00000683543.2; GRCh38) and zebrafish (XM_009296953.3, GRCz11) KMT2D were retrieved from Ensembl and NCBI RefSeq, respectively. The top 7 RBPscan binding motifs were mapped along each transcript and visualized with 5’UTR, CDS, and 3’UTR regions drawn proportionally to scale.

### CellTiter Glo

A375 viability upon DMSO, G3I, or G3Ie treatment was determined using Promega CellTiter-Glo Luminescent Cell Viability Assay (cat. G7570) per manufacturer’s recommendations. Cells were grown to 90% confluency in 96-well microplates (ThermoFisher cat. 136101) and treated with 100 µM sodium arsenite for 4 hours. Media was replaced quickly to remove drug and CellTiter-Glo reagent was added at equal volume for 10 minutes at RT. Luminescence was then recorded using a BioTek Synergy Neo plate reader.

### dsRNA synthesis

*zferv* and *bhikhari-2* LTR dsRNA was generated by *in vitro* transcription upon PCR amplification of each ERV using primers containing a T7 promoter. Each ERV was initially amplified from zebrafish melanoma cDNA using primers without T7 promoter listed in Supp. Table 1. PCR products were purified using Qiagen MinElute PCR Purification Kit (cat. 28006) per manufacturer’s recommended protocol. PCR products were then used as DNA template for a second PCR reaction using the same ERV primers now containing a T7 promoter (Supp. Table 1). PCR products were again purified using Qiagen MinElute PCR Purification Kit. 100 ng of PCR product was then used as template for T7 *in vitro* transcription using MEGAscript T7 Transcription Kit (Invitrogen, cat. AM1334) overnight at 37°C. Synthesized dsRNA was purified with Qiagen RNeasy (cat. 74104) per manufacturer’s recommended protocol. 1000 ng dsRNA was then incubated with Ambion RNase III (Invitrogen, cat. AM2290) for one hour at 37°C. Undigested dsRNAs were treated identically but incubated without enzyme. Digestion was validated by gel electrophoresis using 10% of elutant. The remaining obtained ERV dsRNA was packaged in Lipofectamine 2000 Transfection Reagent (Invitrogen, cat. 11668027) at 1:2 µg:µL ratio in Opti-MEM Reduced Serum Media (Invitrogen, cat. 31985062) and delivered to A375 cells cultured in 24-well glass-bottom plates for 2 hours at 37°C. Mock treatment was used as a negative control and was generated by performing T7 *in vitro* transcription without DNA template and all downstream processing as performed for the ERV dsRNA. HMW poly(I:C) (InvivoGen, cat. tlrl-pic) resuspended to 1 mg/mL in ultrapure H_2_O was used as positive control and delivered at equal ng to ERV dsRNA.

### Statistics

All statistical tests performed are reported in corresponding figure legends. N-values are reported in figure legends or figure panels.

### Data/code availability

All data and code are available upon request.

## Acknowledgements

The authors thank the Zon Lab aquatics staff and Boston Children’s Hospital’s Aquatics Research Program team for fish care and assistance; the Brigham and Women’s Histopathology Core for histology preparation services; the Tissue Biomarker Laboratory at Dana-Farber Cancer Institute for immunostaining human melanomas; Dr. Iwei Yeh for sharing sections of human melanocytic neoplasms; Dr. J. Paul Taylor’s Laboratory for sharing G3I and G3Ie; Dr. Matthew Meyerson for assistance with melanoma mutation genetics; Dr. Kathleen Burns for discussion of ERVs and manuscript review; Dr. David Fisher for discussion of melanoma and manuscript review; Dr. Richard Young for discussion of condensate biology and manuscript review; and all Zon Lab members for their critical feedback and assistance with experimental design, data analysis, and manuscript preparation. This work was supported by funding R01CA103846-14 (LIZ), F32CA294934 (KDD), MoESCS RA 21AG-1F021 (SH, HB, MK), MoESCS RA 25RL-1F024 (SH, HB, MK). JP is funded by a Canadian Institutes for Health Research Postdoctoral Fellowship. RAF is the Bakewell Foundation-Rachleff Innovator of the Damon Runyon Cancer Foundation (DRR-74-23). LIZ is a Howard Hughes Medical Institute Investigator.

## Author Contributions

*Conceptualization* – KDD, LIZ; *Data curation* – KDD, SH, OS, SY; *Formal analysis* – KDD, SH, OS, SY; *Funding acquisition* – KDD, LIZ; *Investigation* – KDD, EF, SH, RAH, ALR, JP, OS; *Methodology* – KDD, SH, OS, JP, RAH; *Project administration* – LIZ; *Resources* – DAK, YZ, MK, DC, RAF; *Software* – SH, OS, SY; *Supervision* – MK, HB, DC, RAF, LIZ; *Validation* – KDD, EF; *Visualization* – KDD, SH, OS; *Writing – original draft* – KDD, LIZ; *Writing – review and editing* – all.

