## Supplementary Figures and Table for "ERV-derived dsRNAs induce stress granules that drive neural crest fate to initiate melanoma"

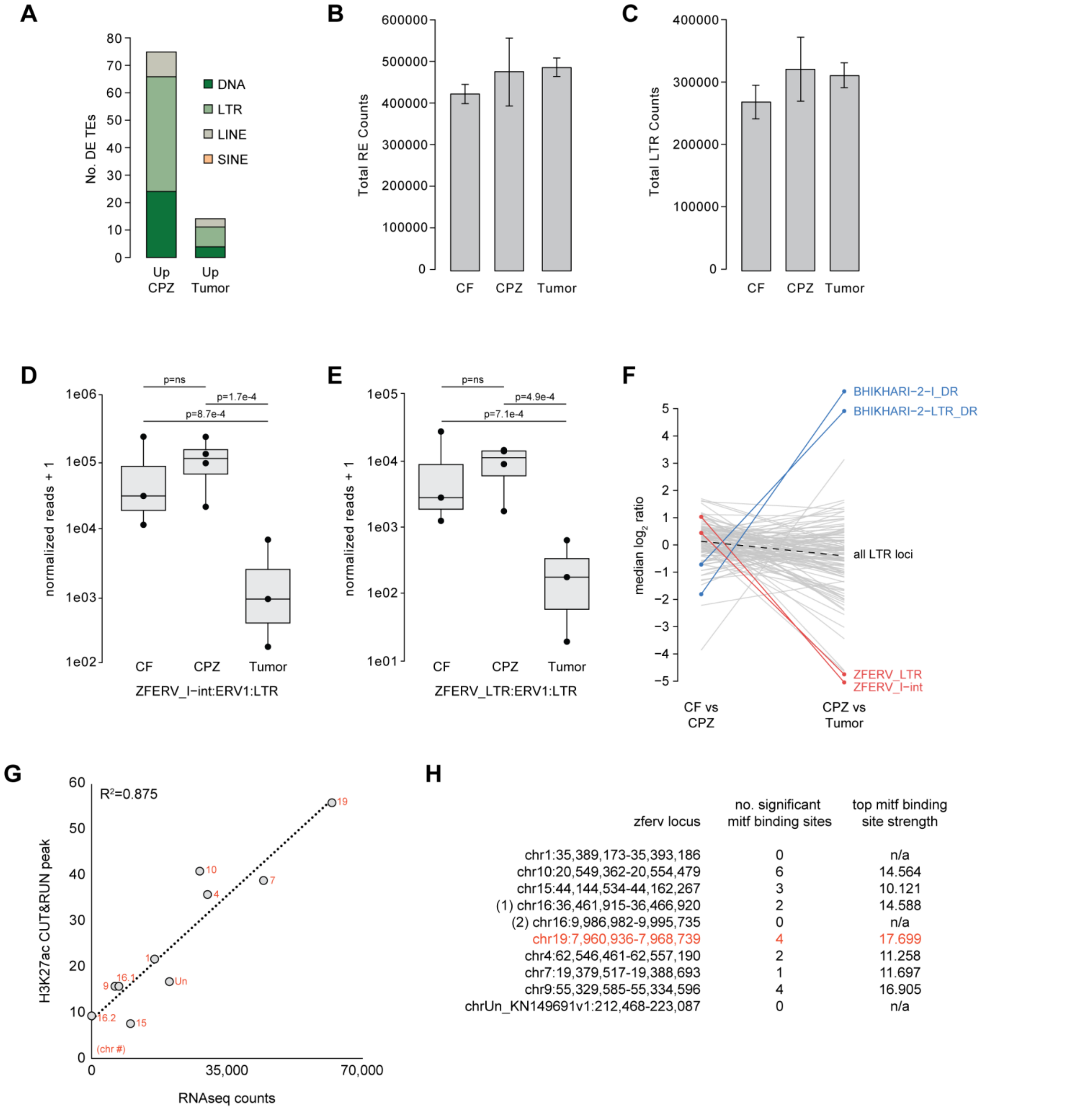


**Supplementary Figure 1.** (**A**) Bar chart showing number of differentially expressed (DE) transposable elements (TEs) in CPZ (n=4) vs tumor (n=3) samples colored by TE class as quantified by TEtranscripts. (**B-C**) Bar charts showing total retroelement (RE; LTR, LINE, SINE) counts (**B**) and LTR counts (**C**) across melanoma stages as quantified by TEtranscripts. Error bars represent SEM. (**D-E**) Box plots showing expression of *zferv* internal (**D**) and LTR (**E**) as quantified and statistically evaluated by SQuIRE. (**F**) Line plot showing median $log_2$ ratio for all *zferv* and *bhikhari-2* loci as quantified by SQuIRE. (**G**) Scatterplot showing correlation between RNAseq read counts (from TEtranscripts) and H3K27ac peak size in CPZs for highest expressed *zferv* loci. (**H**) Table showing number of significant MITF binding sites and top MITF binding site strength +/- 5kb around highest expressed *zferv* loci.

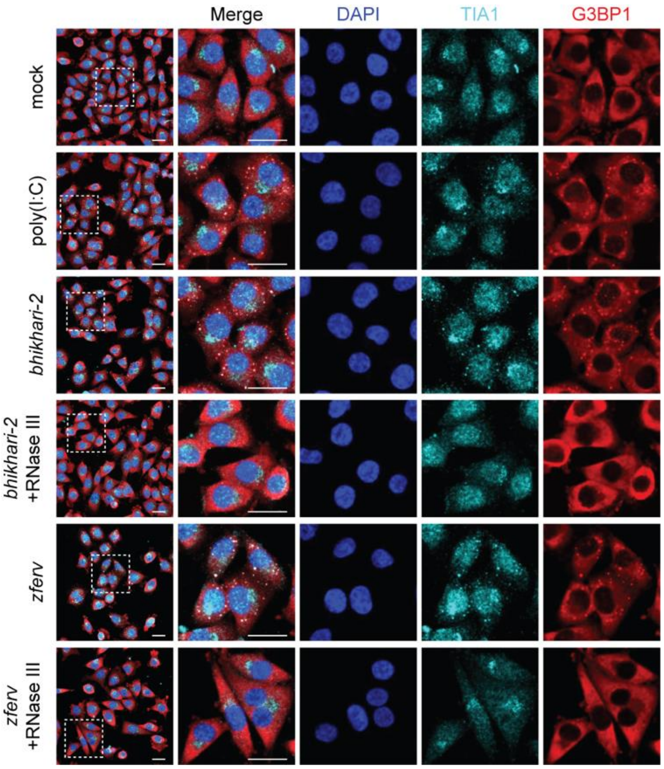


**Supplementary Figure 2.** IF for G3BP1 (red) and TIA1 (cyan) with DAPI counterstain (blue) using A375s lipofected with dsRNA mimic poly(I:C), *bhikhari-2* dsRNA, or *zferv* dsRNA for 2 hours with or without RNase III digestion. Scale bars, 25 µm.

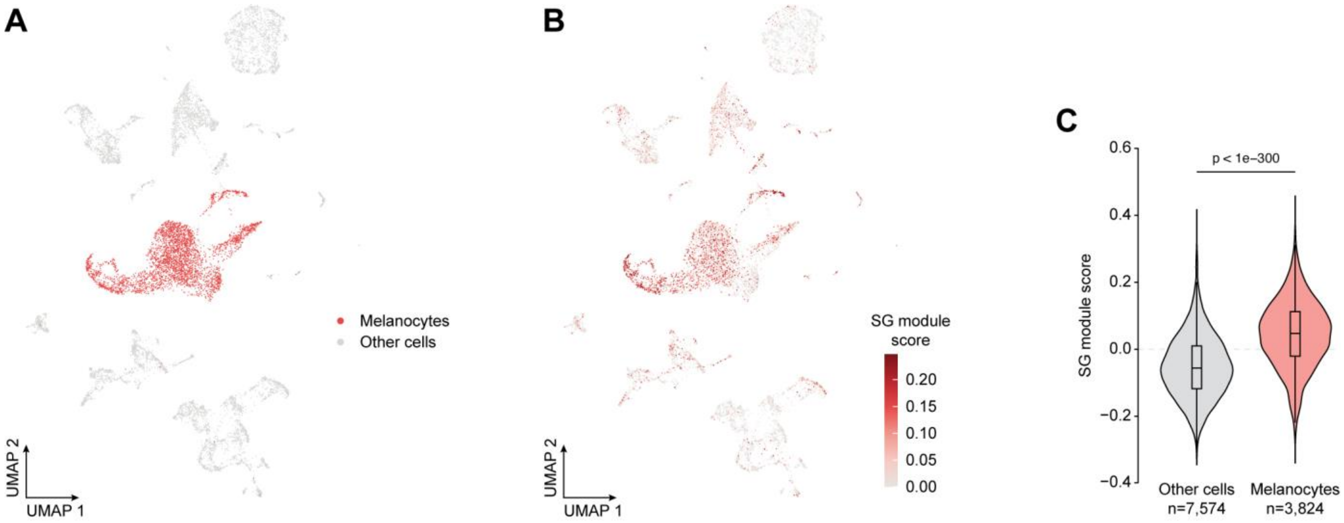


**Supplementary Figure 3.** (**A-C**) UMAP of scRNAseq data from zebrafish melanoma initiation showing (**A**) location of cancerized melanocytes (red) versus captured non-melanocytes (grey) and (**B**) each cell's expression of zebrafish orthologs of core SG genes from Yang et al., 2020 using Seurat's AddModuleScore with (**C**) violin plot of quantification. Statistics determined by Wilcoxon rank-sum test.

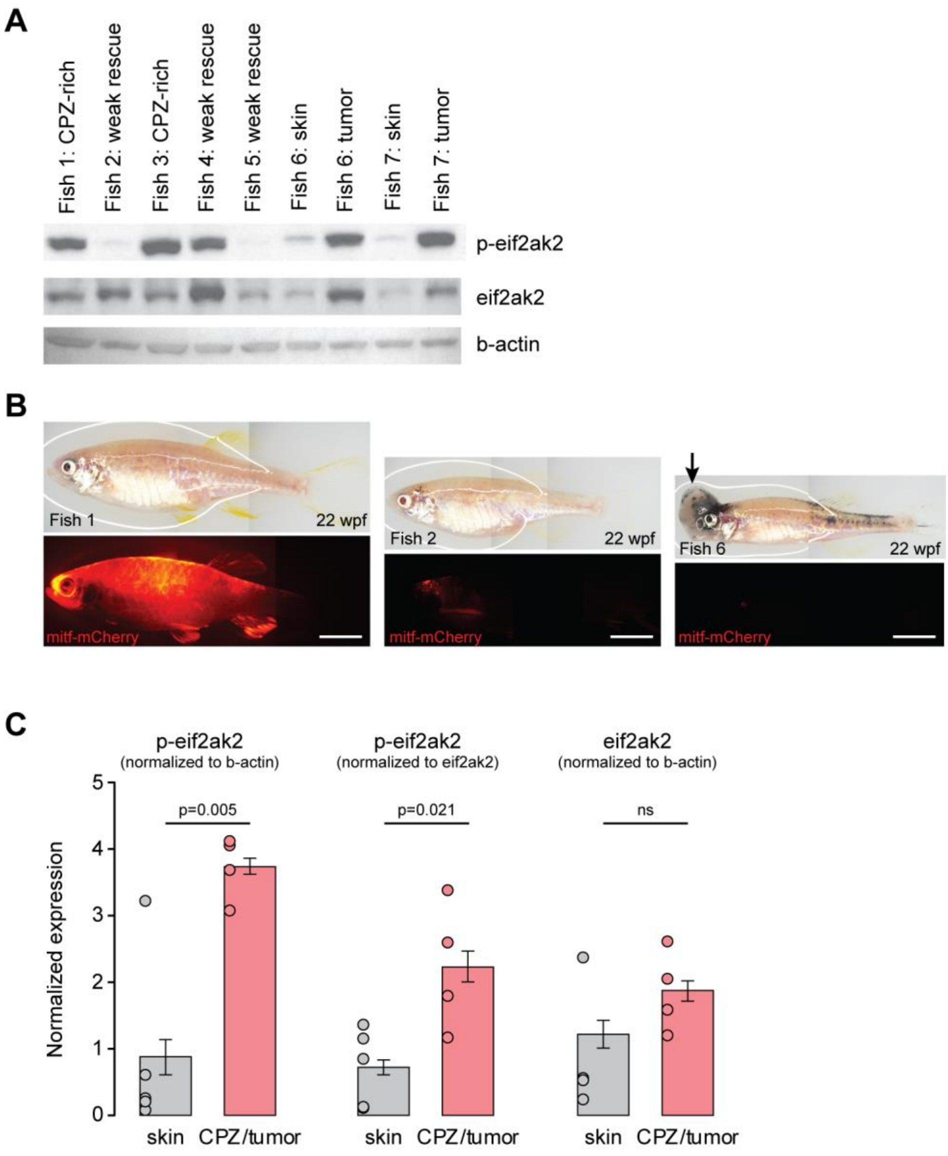


**Supplementary Figure 4.** (**A**) Western blot for p-eif2ak2, eif2ak2, and b-actin from zebrafish skin samples with various levels of melanocyte rescue, CPZ abundance, and tumor status. (**B**) Representative brightfield (top) and fluorescent (bottom) live imaging of zebrafish used for Western blot analysis. (**C**) Bar chart showing quantification of p-eif2ak2 normalized to b-actin (left) or eif2ak2 (middle) and eif2ak2 normalized to b-actin (right). Error bars represent SEM. Significance determined by unpaired T-test. Scale bars, 0.5 cm (**B**).

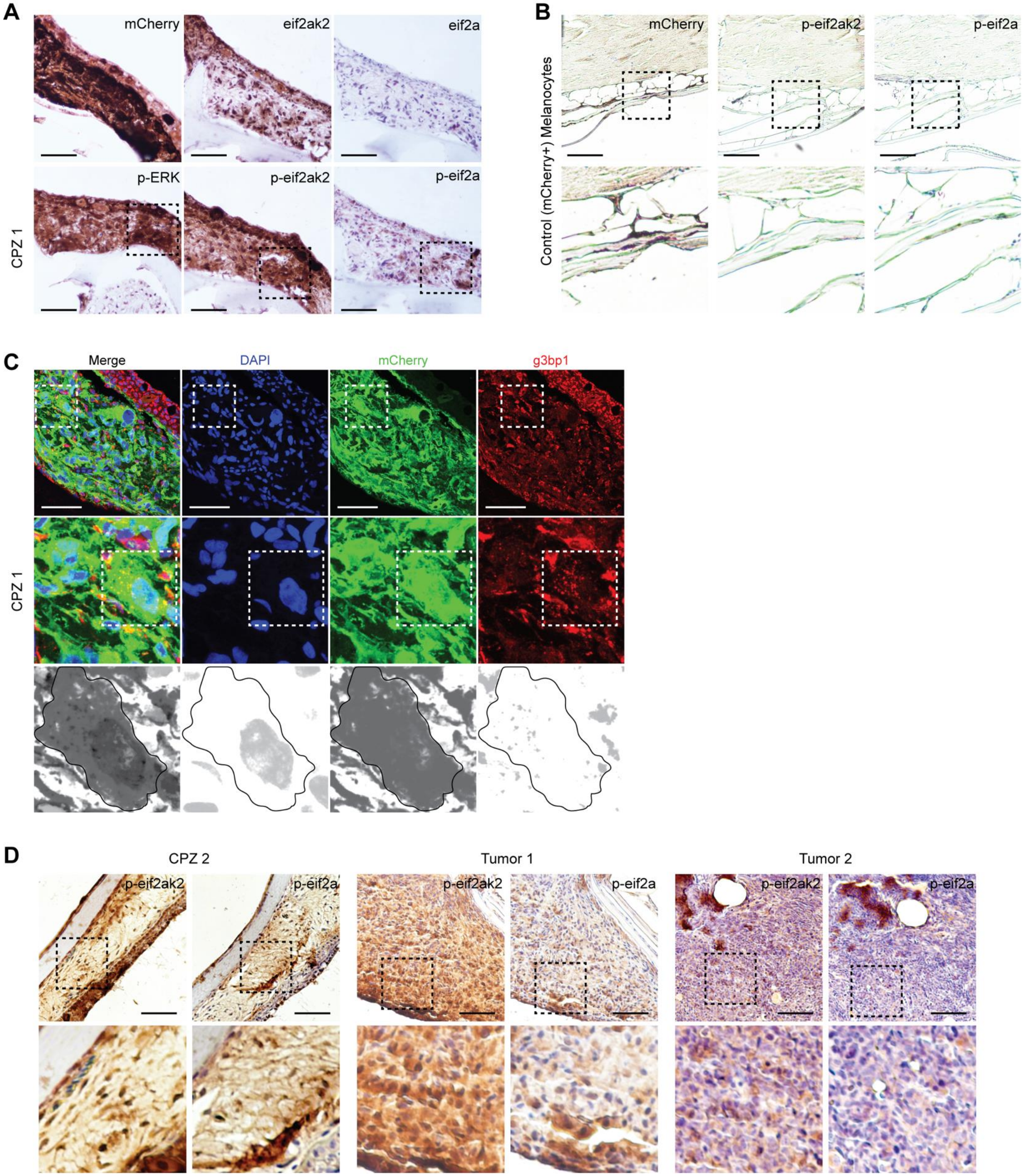


**Supplementary Figure 5.** (**A**) Immunohistochemistry for mCherry, eif2ak2, eif2a, p-ERK, p-eif2ak2, and p-eif2a in representative CPZ. (**B**) Immunohistochemistry for mCherry, p-eif2ak2, and p-eif2a in representative control (mCherry+) melanocytes. (**C**) IF for mCherry (green) and g3bp1 (red) with DAPI nuclear counterstain (blue) in same CPZ stained in (**A**). (**D**) Immunohistochemistry for p-eif2ak2 and p-eif2a in additional CPZ and melanomas. Scale bars, 500 μm (**A**, **B, D**), 50 μm (**C**).

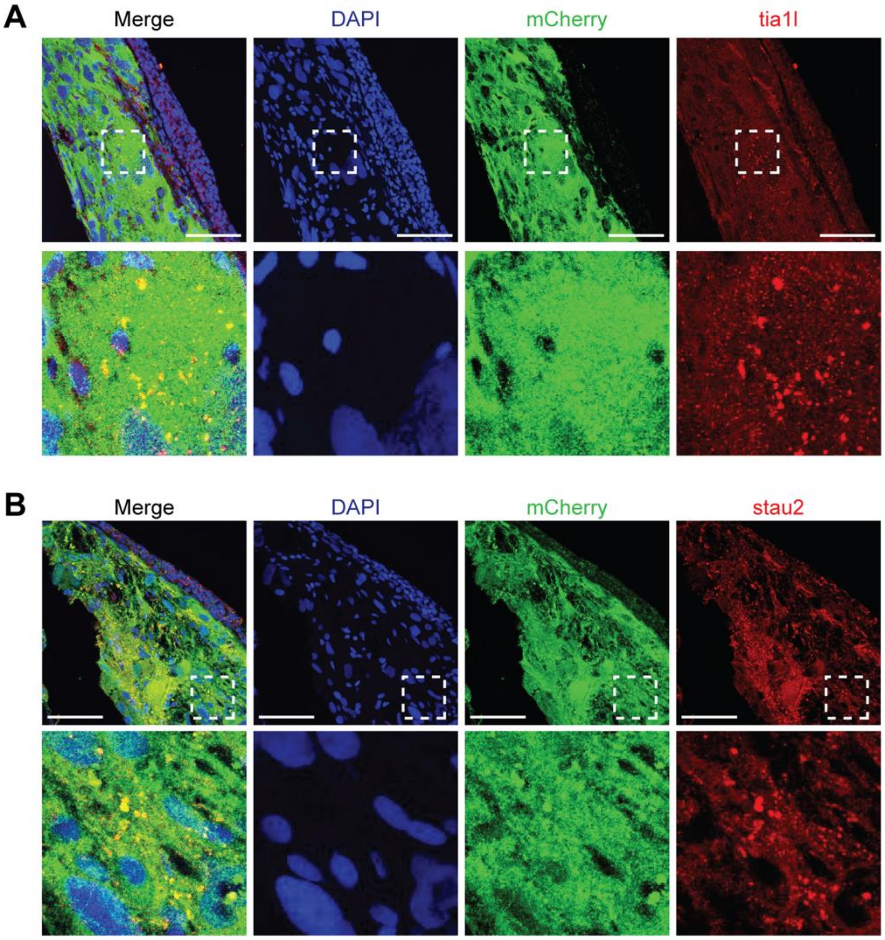


**Supplementary Figure 6.** (**A-B**) IF for mCherry (green) and SG markers (red) tia1l (**A**) and stau2 (**B**) with DAPI nuclear counterstain (blue) in representative CPZ. Scale bars, 50 μm.

**A**

| Class | n | median log2FC | n significant |
|---|---|---|---|
| LTR | 565 | -0.0809 | 20 |
| DNA | 233 | -0.0698 | 2 |
| LINE | 167 | -0.0872 | 2 |
| SINE | 57 | 0.0176 | 1 |
| Unknown | 45 | 0.1325 | 0 |
| Satellite | 21 | 0.0761 | 2 |
| Retroposon | 6 | 0.0795 | 0 |

**B**

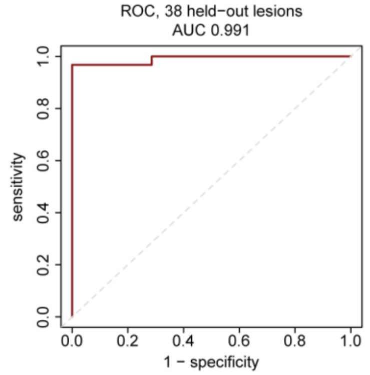


**C** **D**

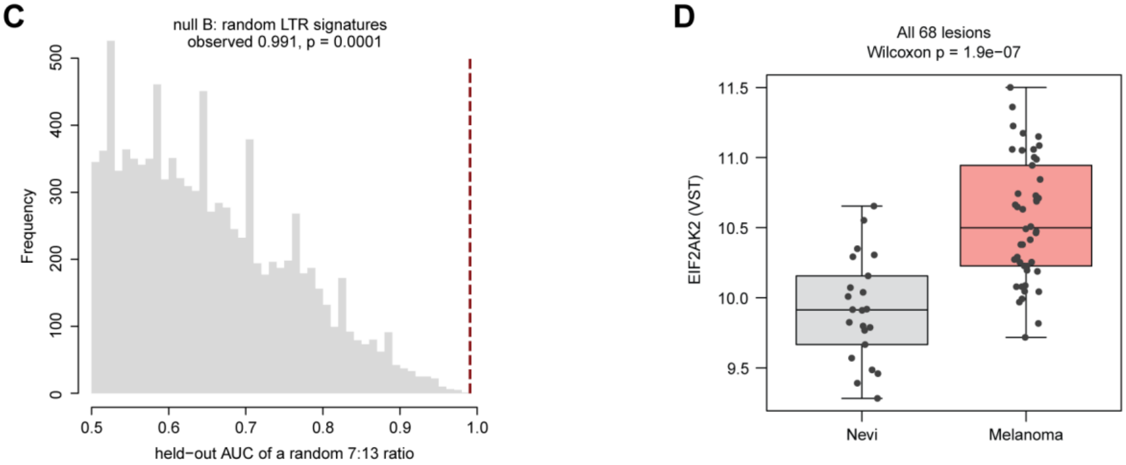


**Supplementary Figure 7.** (**A**) Table showing number (n) of significantly differentially expressed transposable elements in 15 human, paired nevus-melanoma datasets. Significance determined by DESeq2 with a paired design (~ patient + Status); Benjamini-Hochberg-adjusted $p < 0.05$. (**B**) ROC curve showing diagnostic utility of the 20-ERV expression ratio in classifying the unpaired nevi and melanoma datasets. (**C**) Histogram showing AUC frequency upon iterative resampling (10,000 events) of randomly drawn 20-ERV ratio (grey bars) with observed AUC for primary 20-ERV ratio (red dashed line). Random ratios were built from LTR subfamilies outside the signature, matched in number to its 7 and 13 members; empirical $P = (k + 1) / (N + 1) < 0.001$. (**D**) Box plot showing VST-normalized expression of *EIF2AK2* in all 68 analyzed human nevi and melanoma datasets. Significance determined by Wilcoxon rank-sum test.

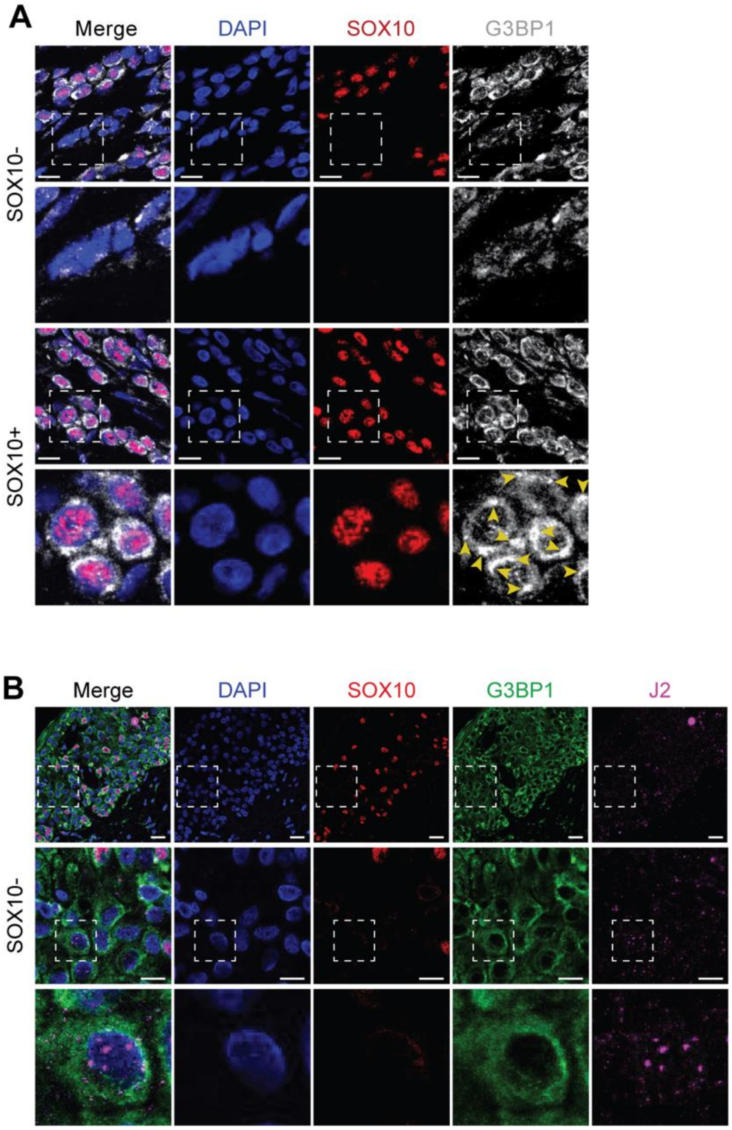


**Supplementary Figure 8.** (**A-B**) IF for G3BP1 and SOX10 (**A**) with dsRNA antibody J2 (**B**) using sections of human nevi. Arrowheads point to SGs. Scale bars, 50 μm.

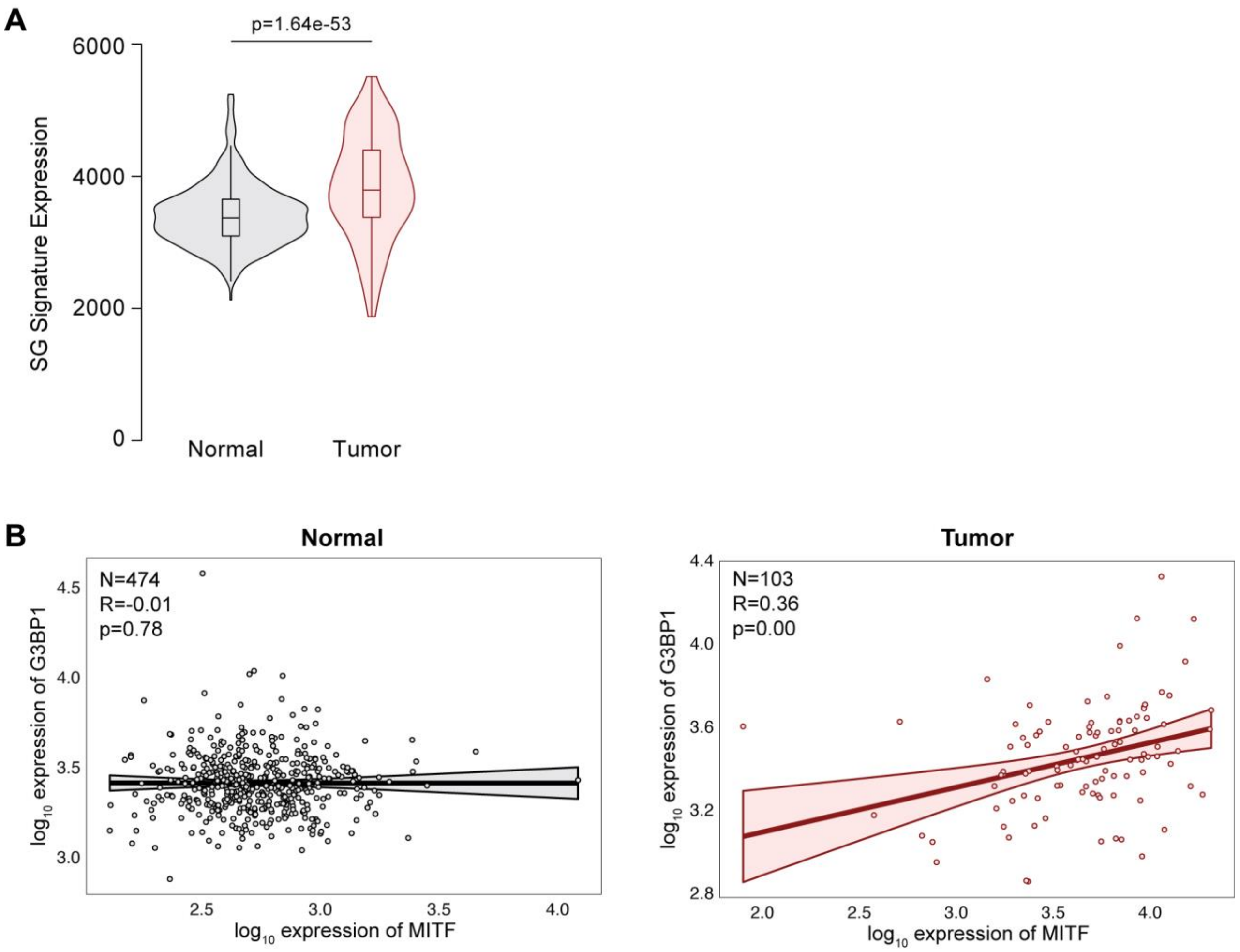


**Supplementary Figure 9.** (**A**) Violin plot showing expression of core SG genes as a signature in human melanoma vs control skin (data and statistics from TNMplot; core SG genes from Yang et al., 2020). (**B**) Scatterplots showing transcriptional correlation of *MITF* and *G3BP1* in normal skin (left) vs human melanoma (right) (data and statistics from TNMplot).

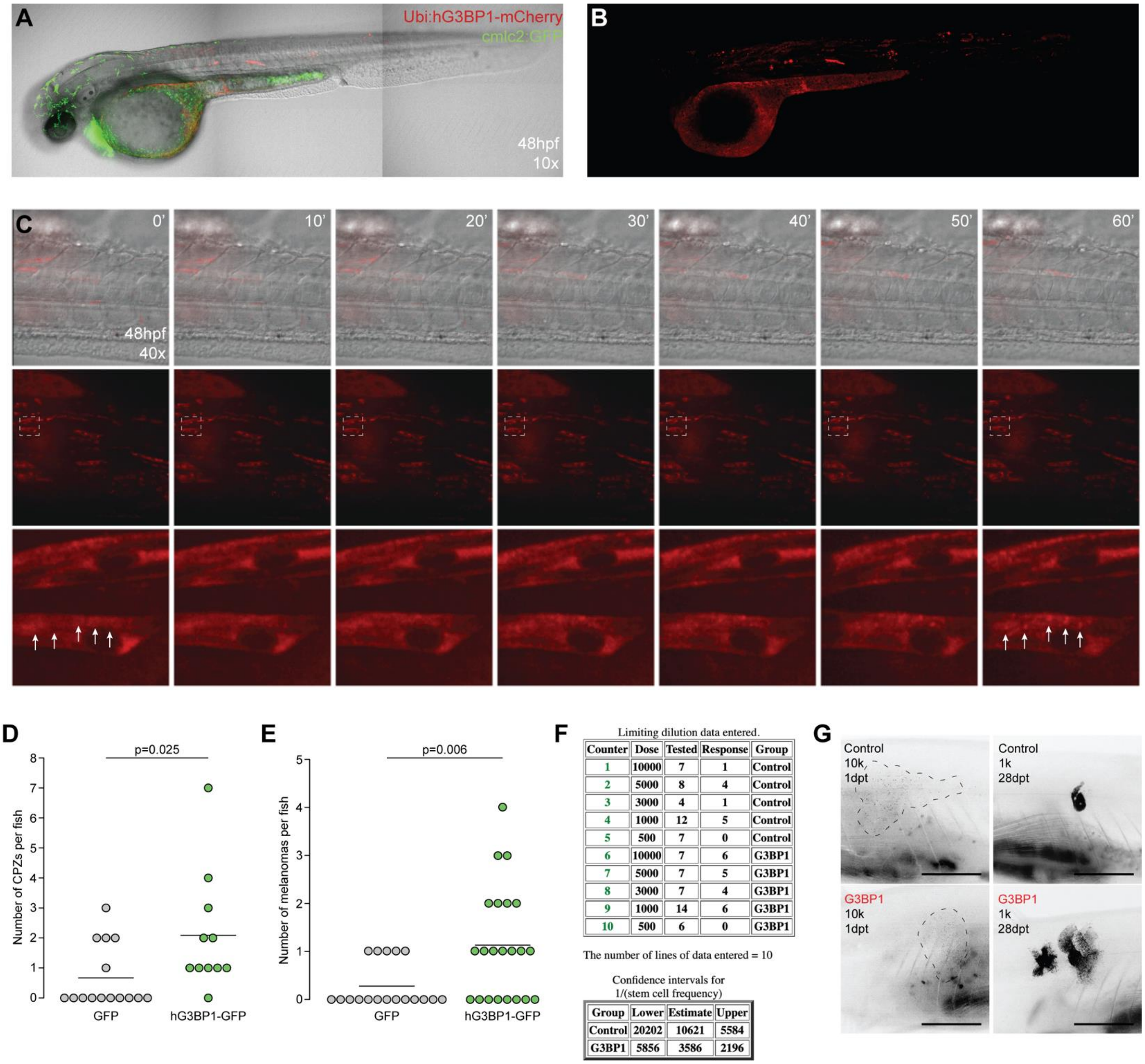


| Counter | Dose | Tested | Response | Group |
|---|---|---|---|---|
| 1 | 10000 | 7 | 1 | Control |
| 2 | 5000 | 8 | 4 | Control |
| 3 | 3000 | 4 | 1 | Control |
| 4 | 1000 | 12 | 5 | Control |
| 5 | 500 | 7 | 0 | Control |
| 6 | 10000 | 7 | 6 | G3BP1 |
| 7 | 5000 | 7 | 5 | G3BP1 |
| 8 | 3000 | 7 | 4 | G3BP1 |
| 9 | 1000 | 14 | 6 | G3BP1 |
| 10 | 500 | 6 | 0 | G3BP1 |

| Group | Lower | Estimate | Upper |
|---|---|---|---|
| Control | 20202 | 10621 | 5584 |
| G3BP1 | 5856 | 3586 | 2196 |

**Supplementary Figure 10.** (**A**) Representative 48-hour post-fertilization (hpf) zebrafish embryo injected with *Ubi:hG3BP1-mCherry* in *pDEST395* (containing green heart reporter *cmlc2:GFP*). (**B**) Representative hG3BP1-mCherry fluorescence of 48hpf zebrafish embryo shown in (**A**). (**C**) Time-lapse of tail region showing hG3BP1-mCherry does not endogenously induce or form SGs at baseline but readily does so upon challenge with a stressor (here, temperature stress of 42°C for 1 hour). Arrowheads show induced SGs. (**D-E**) Quantification of number of CPZs (**D**) and tumors (**E**) in *MCR:GFP* vs *MCR:hG3BP1-GFP* zebrafish. Each dot is one fish. Line represents mean. Significance determined by unpaired T-test. (**F**) Table showing ELDA data generated upon transplant of *MCR:GFP* and *MCR:hG3BP1-GFP* melanoma cells into irradiated casper recipients. (**G**) Representative images of irradiated casper recipients injected with 10k cells at 1-day post-transplant (dpt) (left) and 1k cells at 28 dpt for control (top) and G3BP1-over-expressing (bottom) melanoma cells. Scale bars, 0.5 cm (**G**).

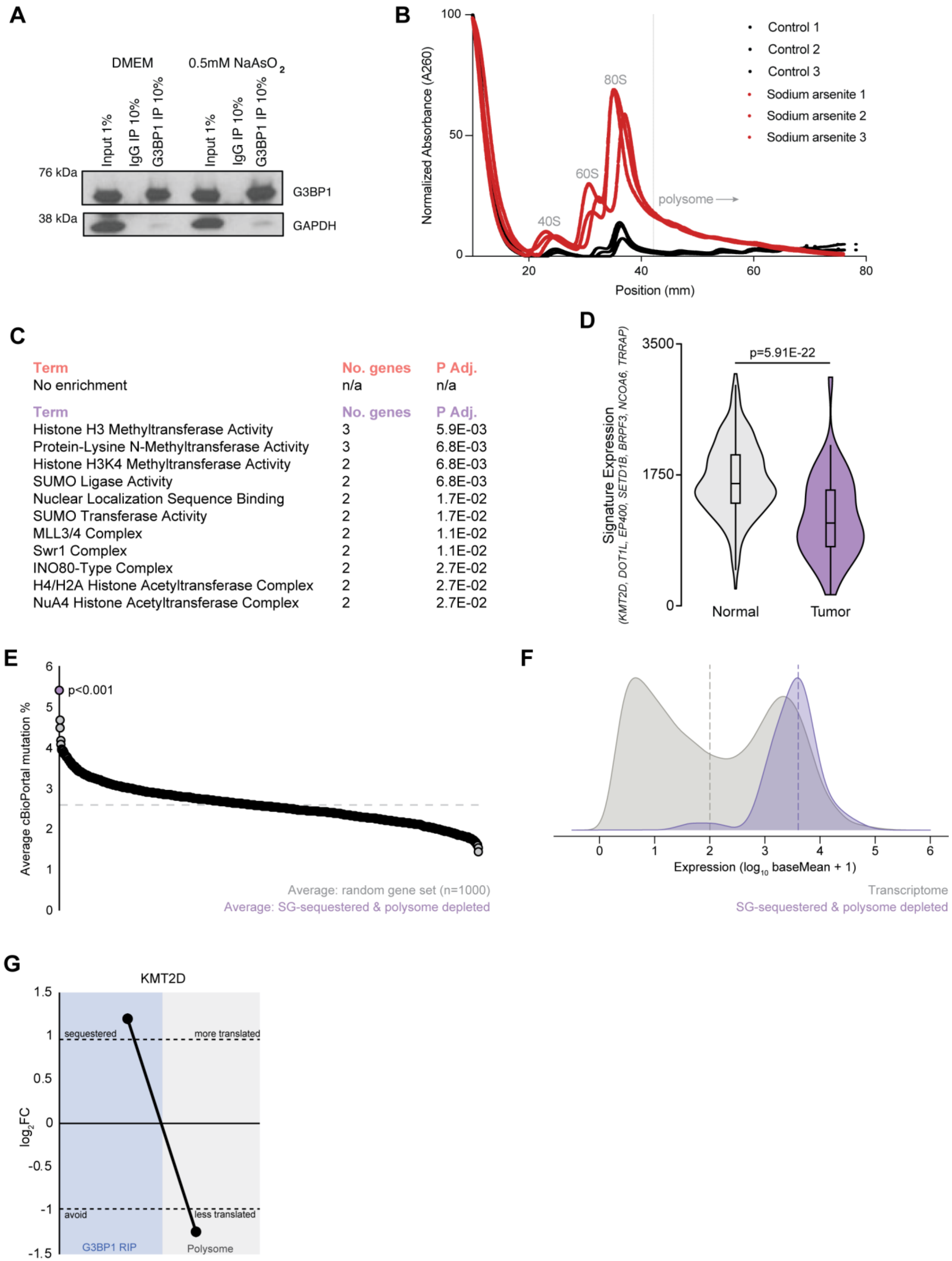


**Supplementary Figure 11.** (**A**) Western blot showing validation of G3BP1-IP from A375 cells used for RIP-seq. (**B**) Sucrose gradient traces for A375 cells stressed with 500 μM sodium arsenite for 1 hour. Dashed line shows threshold past which polysomes were isolated for sequencing. (**C**) Table showing GO-enrichment for genes that avoid G3BP1 and are preferentially translated (top) or sequestered by G3BP1 and less translated (bottom). (**D**) Violin plot showing expression of chromatin-regulating genes identified to be sequestered by G3BP1 and less translated (from cumulative plot) in normal skin vs human melanoma (data and statistics from TNMplot). (**E**) Average cutaneous melanoma mutation rate from iterative sampling of 1000 random sets of 47 genes (size of SG-sequestered and polysome-depleted set) using data curated from cBioPortal. Each dot represents average of one run. Dashed line shows average of all runs. (**F**) Expression distribution of SG-regulated mRNAs (purple) relative to all genes average detected expression in both RIP-seq and polysome-seq inputs (grey). (**G**) Schematic showing $log_2$FC of *KMT2D* in G3BP1 RIP-seq and polysome-seq in A375 cells stressed with 500 μM sodium arsenite for 1 hour.

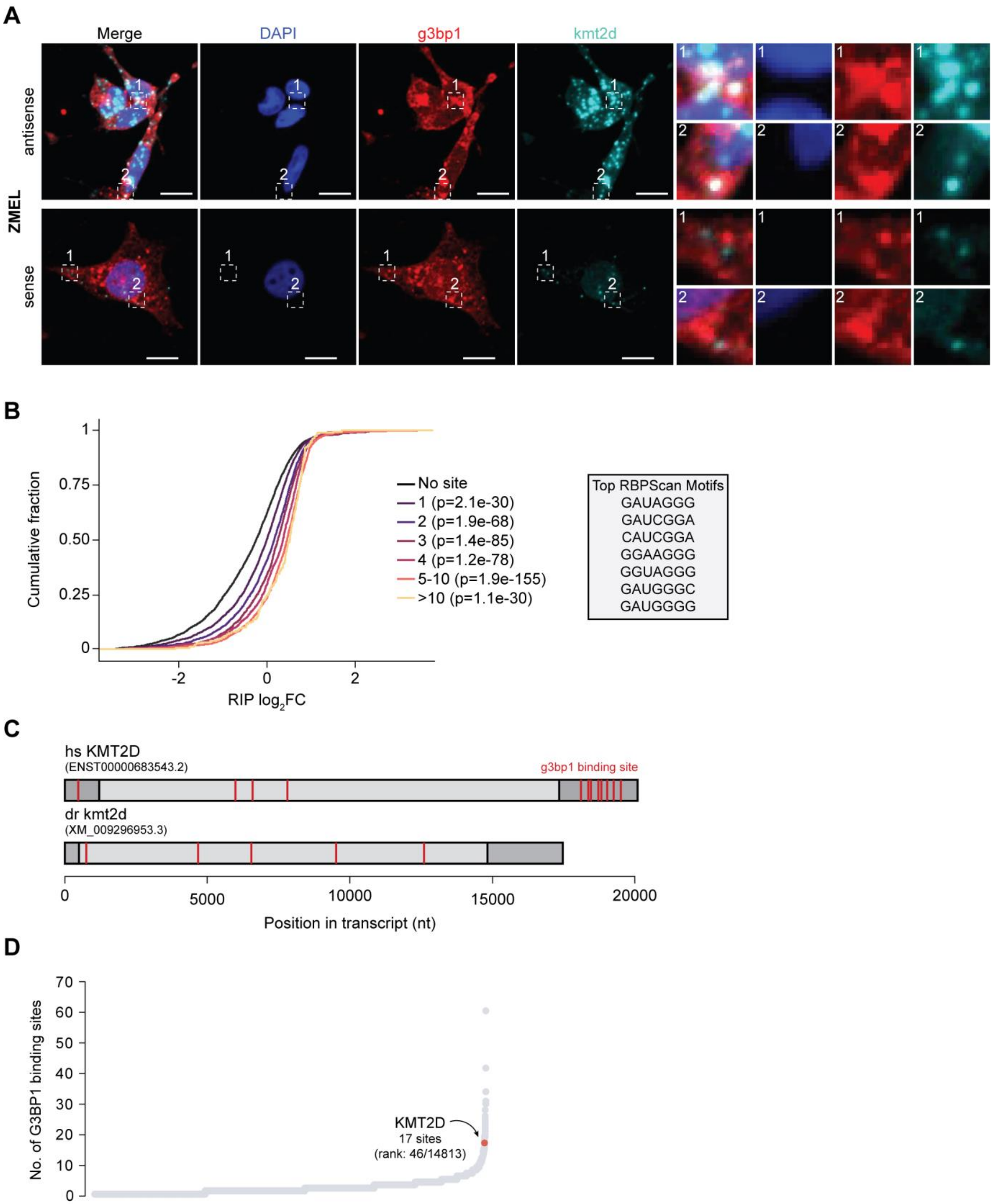


**Supplementary Figure 12.** (**A**) Co-FISH/IF for g3bp1 (IF, red) and *kmt2d* (FISH, cyan) using antisense (top) and sense (bottom) probes with DAPI nuclear counterstain (blue) in ZMEL cells stressed with 500 μM sodium arsenite for 1 hour. (**B**) Traces showing cumulative fraction of genes binned by number of RBPscan-identified G3BP1-binding sites and corresponding G3BP1 RIP-seq $log_2$FC (left) with identified G3BP1 motifs (right). (**C**) Representative transcripts for human *KMT2D* (top) and zebrafish *kmt2d* (bottom) with RBPscan-identified

G3BP1-binding sites (red lines). (**D**) Ordered scatterplot showing number of G3BP1 binding sites per gene expressed in A375 cells. Scale bars, 10 μm (**A**).

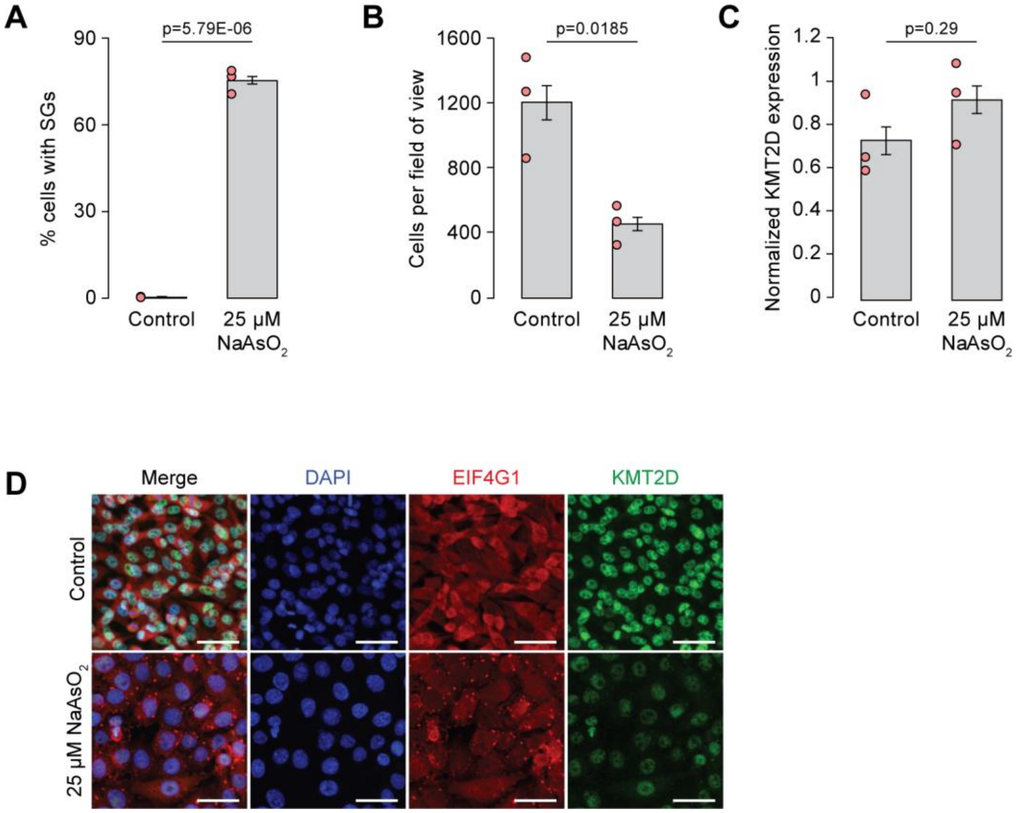


**Supplementary Figure 13.** (**A-B**) Bar chart showing percent of A375 cells with SGs (**A**) and cells per field of view (**B**) upon 24-hour treatment with 25 μM sodium arsenite (n=3). (**C**) Bar chart showing RT-PCR-based quantification of *KMT2D* expression in 24-hour control or 25 μM sodium arsenite-treated A375 cells normalized to GAPDH (n=3). Error bars represent SEM. Statistical significance determined by unpaired T-test. (**D**) IF showing KMT2D (green) downregulation upon chronic SG induction using another SG marker, EIF4G1 (red), with DAPI (blue) nuclear counterstain. Scale bars, 50 μm (**D**).

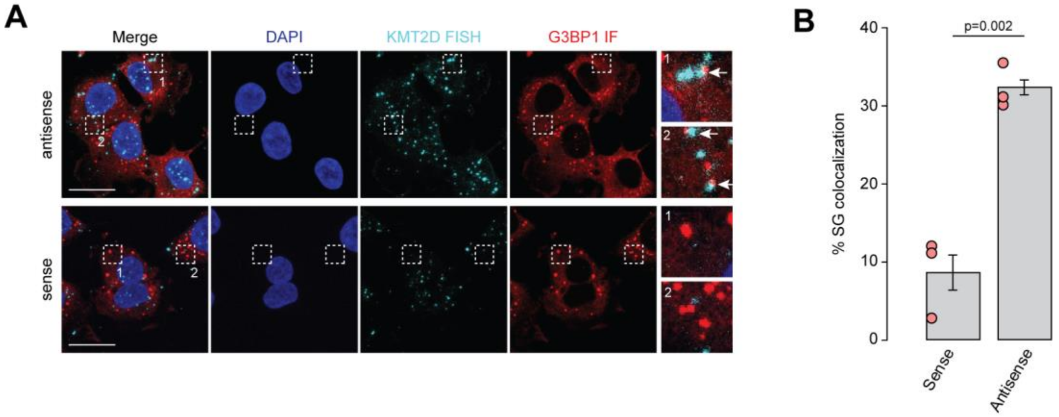


**Supplementary Figure 14.** (**A**) Co-FISH/IF for G3BP1 (IF, red) and *KMT2D* (FISH, cyan) using antisense (top) and sense (bottom) probes with DAPI nuclear counterstain (blue) in A375 cells stressed with 1 μg/mL poly(I:C) for 2 hours. (**B**) Bar chart showing quantification of *KMT2D* FISH puncta colocalization with SGs (n=3). Error bars represent SEM. Significance determined by unpaired T-test. Scale bars, 25 μm (**A**).

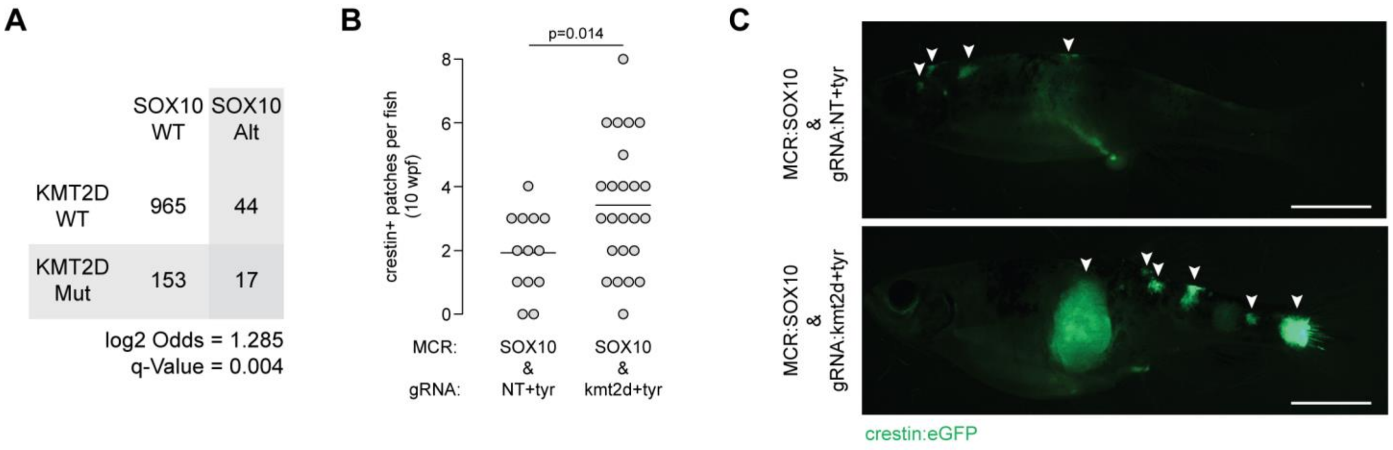


**D**

| Genotype | Tumor | kmt2d gRNA1 indel | kmt2d gRNA2 indel | kmt2d indel sum | kmt2d gRNA1 KO | kmt2d gRNA2 KO | kmt2d KO sum |
|---|---|---|---|---|---|---|---|
| gRNA:NT+tyr & MCR:SOX10 | 1 | 0 | 0 | 0 | 0 | 0 | 0 |
| gRNA:NT+tyr & MCR:SOX10 | 2 | 0 | 0 | 0 | 0 | 0 | 0 |
| gRNA:NT+tyr & MCR:SOX10 | 3 | 0 | 0 | 0 | 0 | 0 | 0 |
| gRNA:NT+tyr & MCR:SOX10 | 4 | 0 | 1 | 1 | 0 | 1 | 1 |
| gRNA:NT+tyr & MCR:SOX10 | 5 | 0 | 0 | 0 | 0 | 0 | 0 |
| gRNA:NT+tyr & MCR:SOX10 | 6 | 0 | 0 | 0 | 0 | 0 | 0 |
| gRNA:NT+tyr & MCR:SOX10 | 7 | 0 | 0 | 0 | 0 | 0 | 0 |
| gRNA:kmt2d+tyr & MCR:SOX10 | 1 | 24 | 0 | 24 | 24 | 0 | 24 |
| gRNA:kmt2d+tyr & MCR:SOX10 | 2 | 36 | 81 | 117 | 36 | 42 | 78 |
| gRNA:kmt2d+tyr & MCR:SOX10 | 3 | 0 | 0 | 0 | 0 | 0 | 0 |
| gRNA:kmt2d+tyr & MCR:SOX10 | 4 | 78 | 91 | 169 | 59 | 91 | 150 |
| gRNA:kmt2d+tyr & MCR:SOX10 | 5 | 37 | 0 | 37 | 19 | 0 | 19 |
| gRNA:kmt2d+tyr & MCR:SOX10 | 6 | 0 | 0 | 0 | 0 | 0 | 0 |
| gRNA:kmt2d+tyr & MCR:SOX10 | 7 | 42 | 2 | 44 | 40 | 2 | 42 |

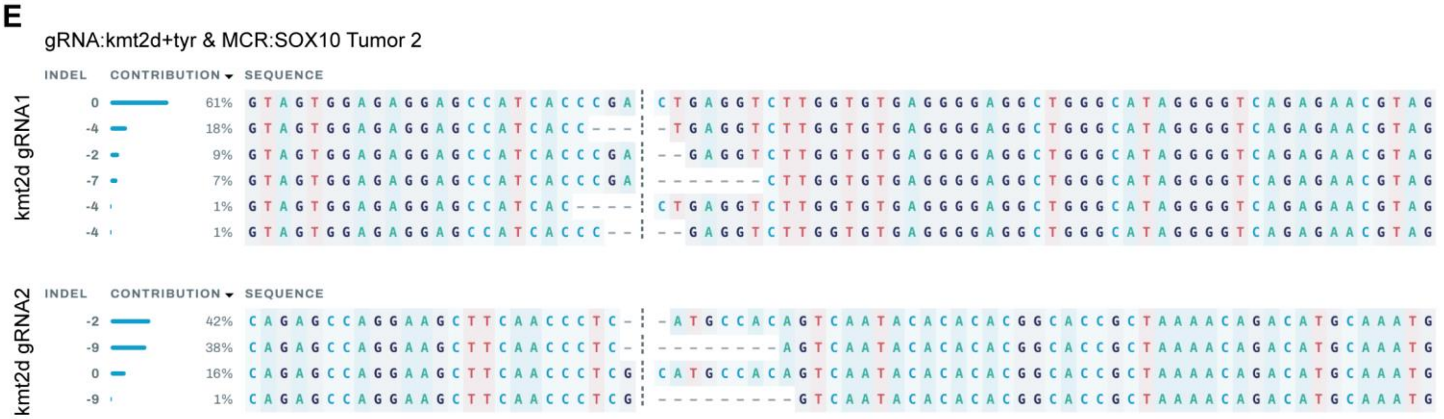


**Supplementary Figure 15.** (**A**) Table showing co-occurrence frequency of *KMT2D* mutation and *SOX10* alteration in human melanoma (data and statistics from cBioPortal). (**B**) Quantification of *crestin*+ patches per *MCR:SOX10 & gRNA:NT+tyr* and *MCR:SOX10 & gRNA:kmt2d+tyr* zebrafish at 10 weeks post-fertilization (wpf). Each dot is one fish. Line represents mean. Significance determined by unpaired T-test. (**C**) Representative fluorescent live imaging of crestin-eGFP patches in 10wpf *MCR:SOX10 & gRNA:NT+tyr* (top) and *MCR:SOX10 & gRNA:kmt2d+tyr* (bottom) zebrafish. (**D**) Table showing indel and predicted knockout (KO) editing of *kmt2d* at the loci targeted by *kmt2d* gRNA1 and *kmt2d* gRNA2. (**E**) Editing analysis of *kmt2d* gRNA1 (top) and *kmt2d* gRNA2 (bottom) for a representative *MCR:SOX10 & gRNA:kmt2d+tyr* tumor, as generated by Synthego ICE. Scale bars, 0.5 cm (**C**).

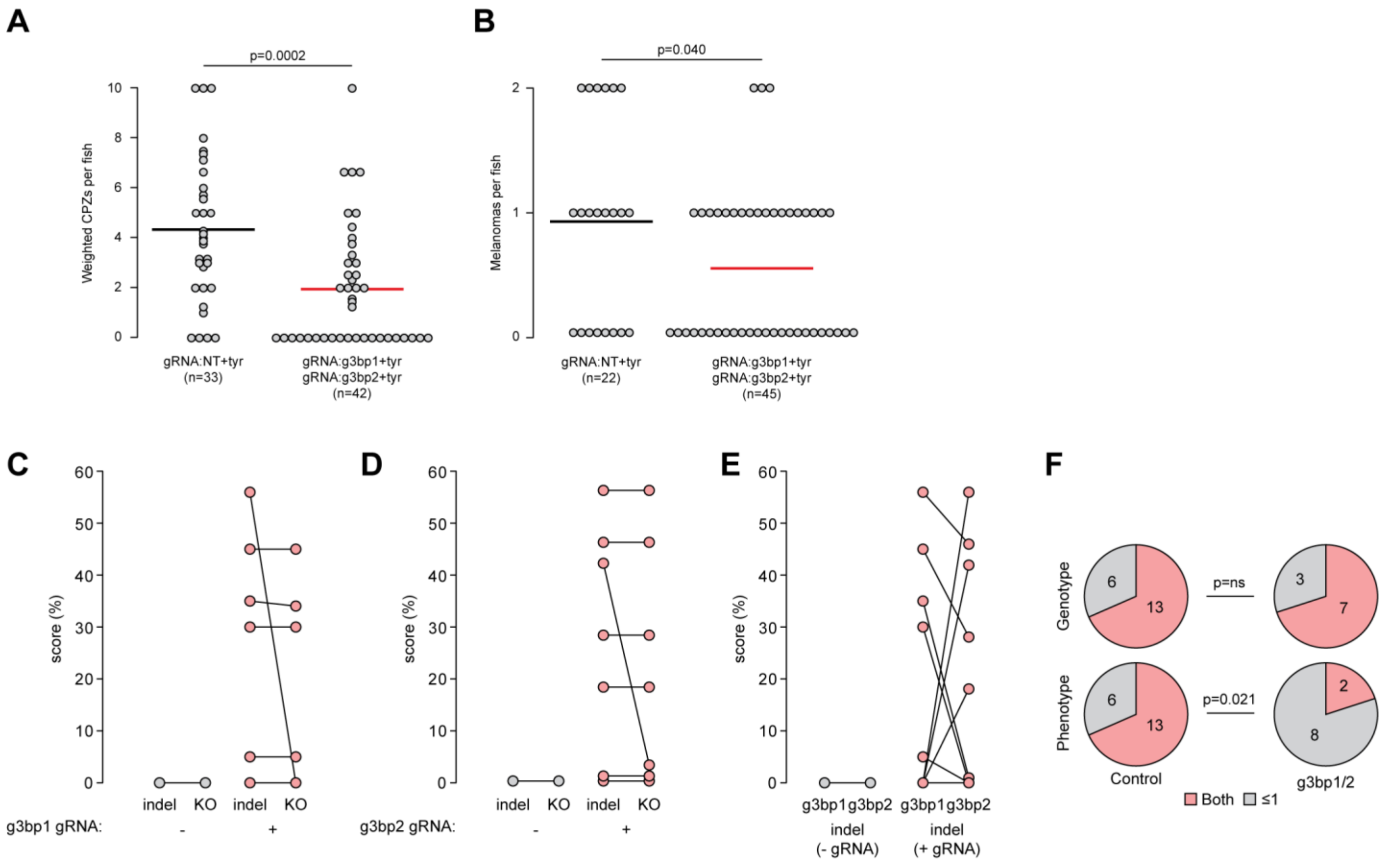


G

| Genotype | Tumor | g3bp1 indel | g3bp2 indel | indel sum | g3bp1 KO | g3bp2 KO | KO sum | detect g3bp1 and g3bp2 plasmid? |
|---|---|---|---|---|---|---|---|---|
| gRNA:NT+tyr | 1 | 0 | 0 | 0 | 0 | 0 | 0 | n/a |
| gRNA:NT+tyr | 2 | 0 | 0 | 0 | 0 | 0 | 0 | n/a |
| gRNA:NT+tyr | 3 | 0 | 0 | 0 | 0 | 0 | 0 | n/a |
| gRNA:NT+tyr | 4 | 0 | 0 | 0 | 0 | 0 | 0 | n/a |
| gRNA:g3bp1/2+tyr | 1 | 45 | 28 | 73 | 45 | 28 | 73 | both |
| gRNA:g3bp1/2+tyr | 2 | 0 | 18 | 18 | 0 | 18 | 18 | both |
| gRNA:g3bp1/2+tyr | 3 | 35 | 1 | 36 | 34 | 1 | 35 | both |
| gRNA:g3bp1/2+tyr | 4 | 0 | 0 | 0 | 0 | 0 | 0 | g3bp2 only |
| gRNA:g3bp1/2+tyr | 5 | 5 | 0 | 5 | 5 | 0 | 5 | both |
| gRNA:g3bp1/2+tyr | 6 | 30 | 0 | 30 | 30 | 0 | 30 | both |
| gRNA:g3bp1/2+tyr | 7 | 56 | 46 | 102 | 0 | 46 | 46 | both |
| gRNA:g3bp1/2+tyr | 8 | 0 | 42 | 42 | 0 | 3 | 3 | both |
| gRNA:g3bp1/2+tyr | 9 | 0 | 56 | 56 | 0 | 56 | 56 | both |
| gRNA:g3bp1/2+tyr | 10 | 0 | 0 | 0 | 0 | 0 | 0 | g3bp2 only |

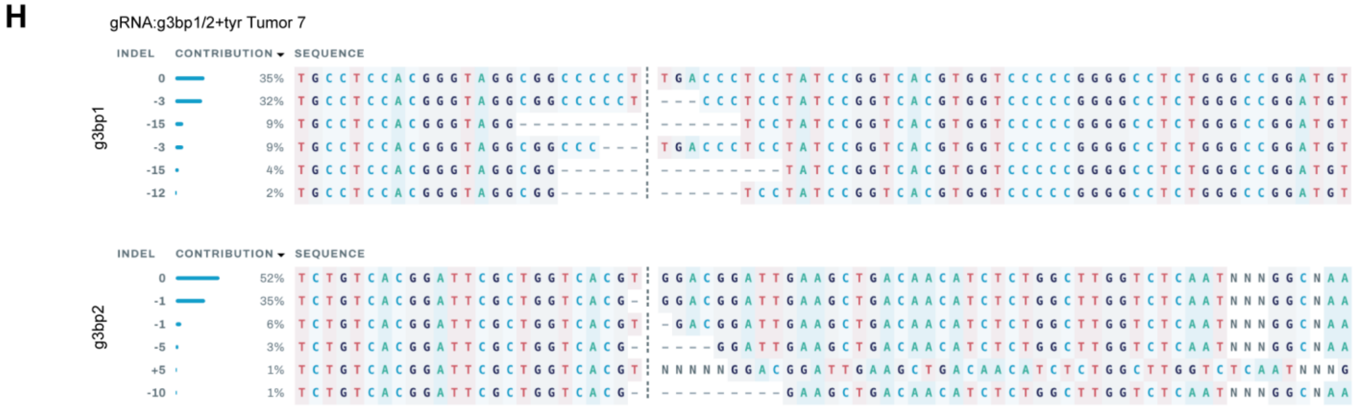


**Supplementary Figure 16.** (**A-B**) Quantification of number of CPZs (**A**) and melanomas (**B**) in control (*gRNA:NT+tyr*) and *g3bp1/2-KO* (*gRNA:g3bp1+tyr & gRNA:g3bp2+tyr*) zebrafish. Each dot is one fish. Line represents mean. Significance determined by unpaired T-test. (**C-E**) Percent indel and predicted knockout (KO) score for *g3bp1* (**C**), *g3bp2* (**D**), and combined (**E**) in control and *g3bp1/2-KO* zebrafish melanomas. (**F**) Pie chart showing PCR-based detection of plasmids in control (*MCR:GFP* + *MCR:mCherry*) and *g3bp1/2-KO* melanoma (top) with observation rate of expressed phenotype in control (fluorophore detection) and *g3bp1/2-KO* (genomic editing) melanomas (bottom). Significance determined by Fisher's exact test. (**G**) Table showing *g3bp1* and *g3bp2* indel and predicted KO percentages in control and *g3bp1/2-KO* melanomas. (**H**) Editing analysis of *g3bp1* (top) and *g3bp2* (bottom) for a representative, co-edited *g3bp1/2-KO* zebrafish displaying maintenance of one in-frame allele (*g3bp1*), as generated by Synthego ICE.

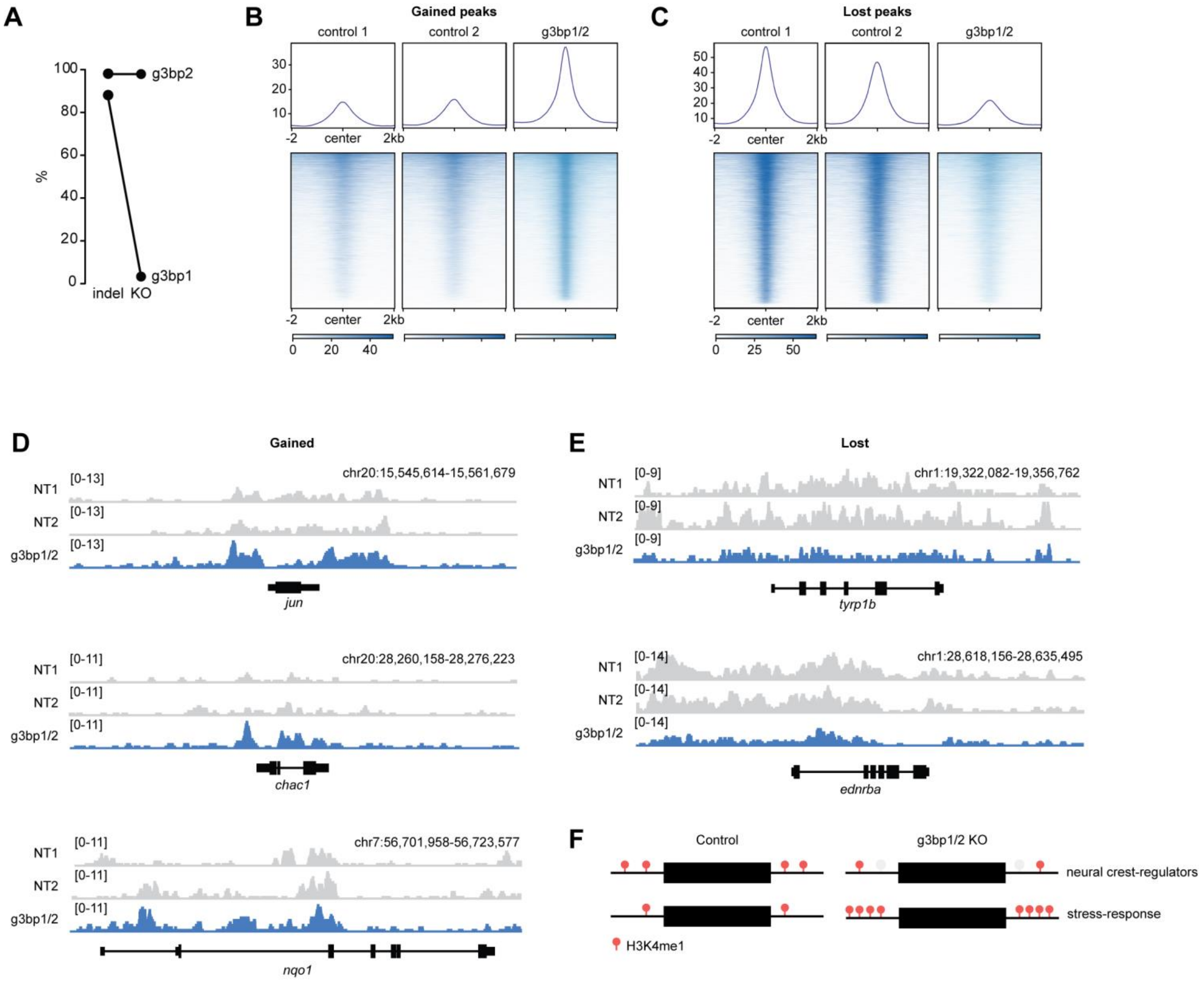


**Supplementary Figure 17.** (**A**) Percent indel and predicted knockout (KO) for *g3bp1* and *g3bp2* in *g3bp1/2-KO* (*gRNA:g3bp1+tyr & gRNA:g3bp2+tyr*) zebrafish melanoma harvested for H3K4me1 CUT&RUN. (**B-C**) Heatmaps showing gained (**B**) and lost (**C**) H3K4me1 peaks in *g3bp1/2-KO* melanoma +/-2kb from center. (**D-E**) Representative IGV tracks for gained (**D**) and lost (**E**) H3K4me1 peaks in *g3bp1/2-KO* melanoma. (**F**) Model showing H3K4me1 redistribution in *g3bp1/2-KO* melanoma.

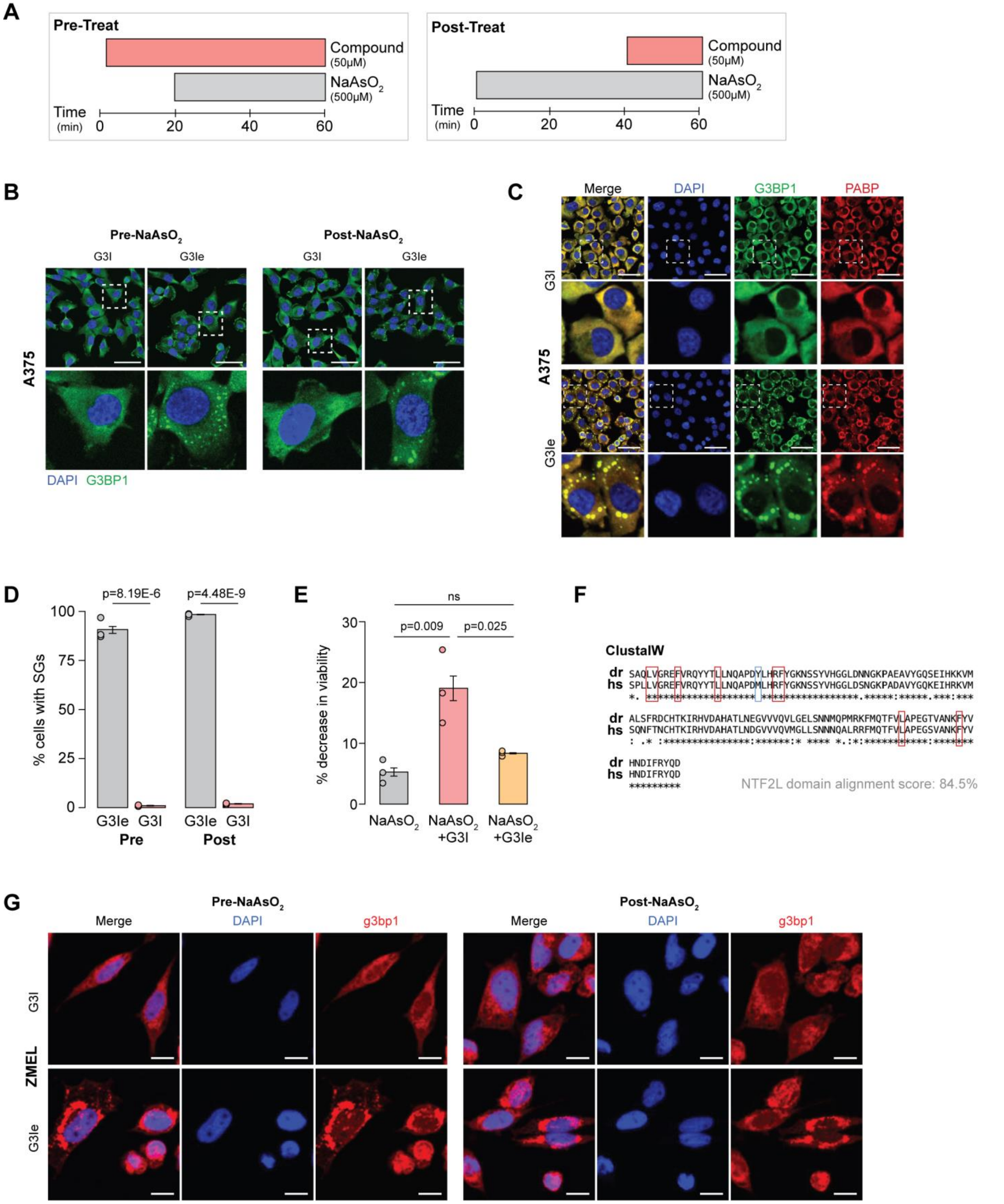


**Supplementary Figure 18.** (**A**) Schematic depicting G3I/G3Ie incubation paradigm prior to (left, pre-treat) or after (right, post-treat) 500 µM sodium arsenite for A375 and ZMEL cells. (**B**) IF for G3BP1 (green) with DAPI nuclear counterstain (blue) on A375 cells treated with 500 µM sodium arsenite and pre- or post-treated with 50 µM G3I or G3Ie. (**C**) IF for G3BP1 (green) and secondary SG marker PABP (red) in A375 cells post-treated with

G3I (top) or G3Ie (bottom) shows complete SG dissolution beyond G3BP1 inhibition. (**D**) Bar chart showing percent of A375 cells containing SGs when pre- or post-treated with G3Ie or G3I and exposed to 500 µM sodium arsenite (n=3). Error bars represent SEM. Significance determined by unpaired T-test. (**E**) Bar chart showing CellTiter Glo Assay of A375 cells treated with 100 µM sodium arsenite for 4 hours with no drug, G3I, or G3Ie (n=3). Error bars represent SEM. Significance determined by ANOVA with post-hoc Tukey's test. (**F**) ClustalW alignment of zebrafish (top) and human (bottom) G3BP1 NTF2L domain. Boxed amino acids show G3I targets. (**G**) IF for g3bp1 (red) with DAPI nuclear counterstain (blue) on ZMEL cells stressed with 500 µM sodium arsenite and pre- or post-treated with 50 µM G3I or G3Ie. Scale bars, 50 µm (**B**, **C**), 10 µm (**G**).

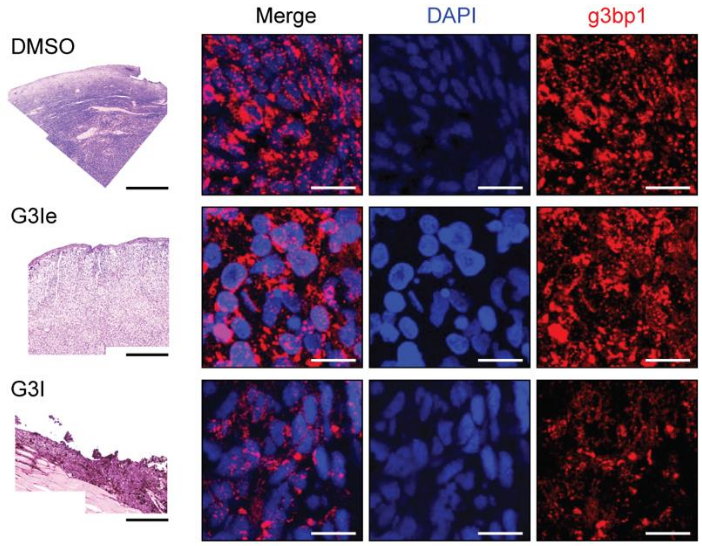


**Supplementary Figure 19.** H&E (left) and IF (right) for g3bp1 (red) with DAPI nuclear counterstain (blue) for representative DMSO (top), G3Ie (middle), or G3I (bottom) injected melanomas. Scale bars, 500 μm (H&E), 10 μm (IF).

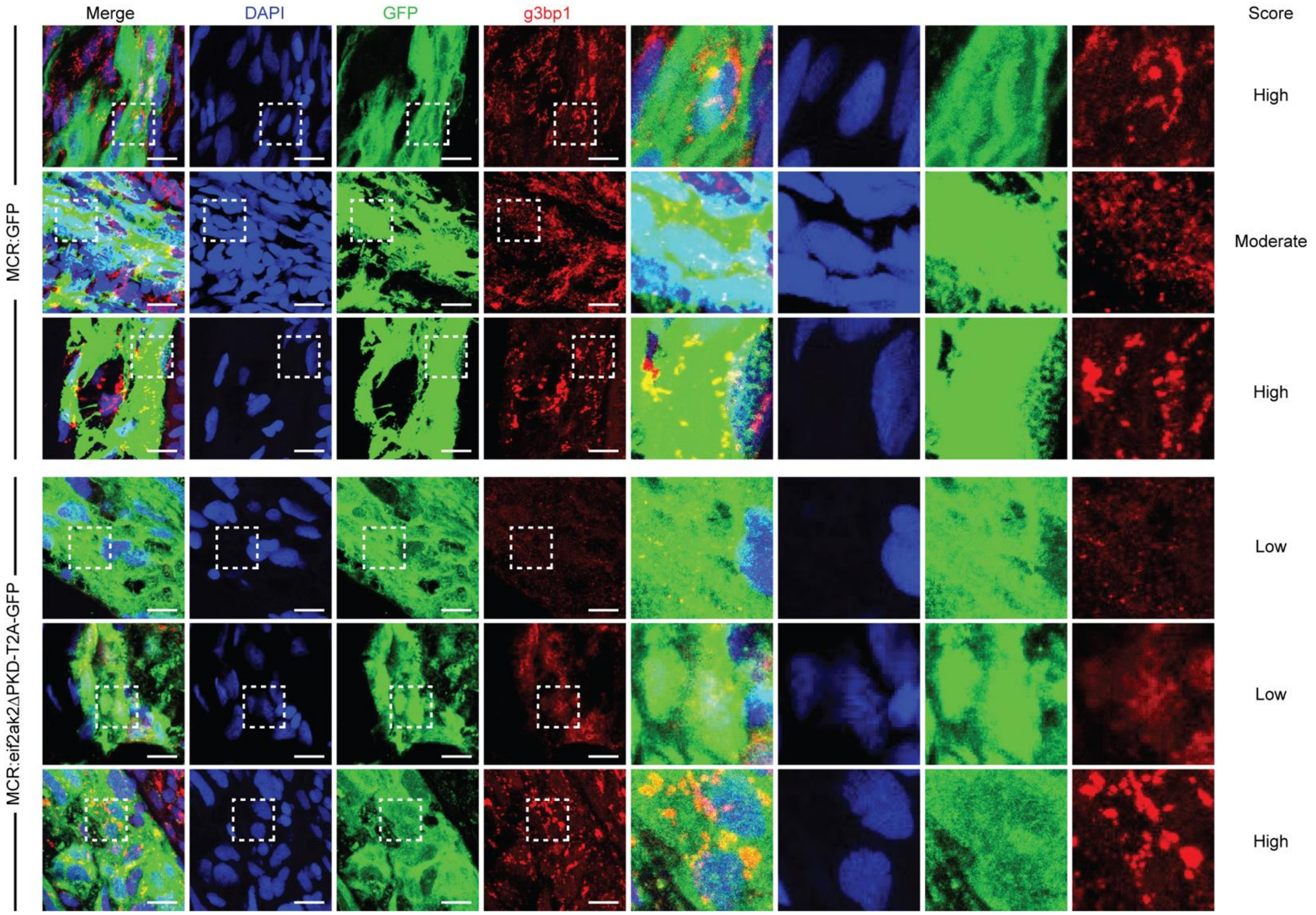


**Supplementary Figure 20.** IF for GFP (green) and g3bp1 (red) with DAPI nuclear counterstain (blue) showing representative SG scoring in *MCR:GFP* (top) or *MCR:eif2ak2ΔPKD-T2A-GFP* (bottom) CPZs. Scale bars, 10 µm.

**Supplementary Table 1.** PCR primers used in this study.

| Gene | Species | Purpose |
|---|---|---|
| *g3bp1* | *Zebrafish* | *Genotyping (CRISPR editing)* |
| Fwd: 5'-GTAGAGGAAAAGAAGACCCGCT-3' | | |
| Rvs: 5'-TTGTGTTAACCAGGGTGACTTG-3' | | |
| *g3bp2* | *Zebrafish* | *Genotyping (CRISPR editing)* |
| Fwd: 5'-ATTAGCCCAGAGTTGAGACCAA-3' | | |
| Rvs: 5'-AAAACATTTCAGATCAACTGCC-3' | | |
| *kmt2d* | *Zebrafish* | *Genotyping (CRISPR editing, gRNA 1)* |
| Fwd: 5'-AGTTACCGCCAATCTCCATCTA-3' | | |
| Rvs: 5'-CACAGAGAGGACATCATTGGAA-3' | | |
| *kmt2d* | *Zebrafish* | *Genotyping (CRISPR editing, gRNA 2)* |
| Fwd: 5'-AGCTTGTACAGAAGTTTGGCAA-3' | | |
| Rvs: 5'-GGAAAAGTACACTTTAGAAAACAGC-3' | | |
| *zferv* | *Zebrafish* | *dsRNA synthesis* |
| Fwd: 5'-CACCTGAGGGAATTATATTTTTGAGGG-3' | | |
| Rvs: 5'-TGTTGGGAATGGTTTTTCCCTCCC-3' | | |
| *bhikhari-2* | *Zebrafish* | *dsRNA synthesis* |
| Fwd: 5'-CACCTGAAGGAACTCATCGGCATA-3' | | |
| Rvs: 5'-TGAGTACAAAGATTTATTGAGCTAC-3' | | |
| *zferv* | *Zebrafish* | *dsRNA synthesis (with T7)* |
| Fwd: 5'-CAGTGAATTGTAATACGACTCACTATAGGATGGGTGAGGGAATTATATTTTTGAGGG-3' | | |
| Rvs: 5'-CAGTGAATTGTAATACGACTCACTATAGGATGGGTGTTGGGAATGGTTTTTCCCTCCC-3' | | |
| *bhikhari-2* | *Zebrafish* | *dsRNA synthesis (with T7)* |
| Fwd: 5'-CAGTGAATTGTAATACGACTCACTATAGGATGGGTGAAGGAACTCATCGGCATA-3' | | |
| Rvs: 5'-CAGTGAATTGTAATACGACTCACTATAGGATGGGTGAGTACAAAGATTTATTGAGCTAC-3' | | |
| *KMT2D* | *Human* | *RT-PCR* |
| Fwd: 5'-GGAATGGGTAGCTCTTTGGCGA-3' | | |
| Rvs: 5'-TGCCGAATCAGCAGCTCTCGTA-3' | | |
| *GAPDH* | *Human* | *RT-PCR* |
| Fwd: 5'-CAACGAATTTGGCTACAGCAACAGG-3' | | |
| Rvs: 5'-GGCAACTGTGAGGAGGGGAGATT-3' | | |
| *KMT2D* | *Human* | *In situ probe* |
| Fwd: 5'-CAGTGAATTGTAATACGACTCACTATAGGAATGGGTAGCTCTTTGGCGA-3' | | |
| Rvs: 5'-CAGTGAATTGTAATACGACTCACTATAGGTGCCGAATCAGCAGCTCTCGTA-3' | | |
